# Neurovascular unit alterations in the growth restricted newborn are improved following ibuprofen treatment

**DOI:** 10.1101/2021.02.15.431329

**Authors:** Kirat K. Chand, Stephanie M. Miller, Gary J. Cowin, Lipsa Mohanty, Jany Pienaar, Paul B. Colditz, Stella Tracey Bjorkman, Julie A. Wixey

## Abstract

Fetal brain development is particularly vulnerable to the effects of fetal growth restriction (FGR) and abnormal neurodevelopment is common in the FGR infant. Adverse outcomes range from behavioural and learning disorders through to cerebral palsy. Unfortunately, no treatment exists to protect the FGR newborn brain. Recent evidence suggests inflammation may play a key role in the mechanism responsible for the progression of brain impairment in the FGR newborn, including disruption to the neurovascular unit (NVU). We explored whether ibuprofen, a non-steroidal anti-inflammatory drug, could reduce NVU disruption and brain impairment in the FGR newborn. We used a preclinical piglet model of growth restriction in which FGR occurs spontaneously. Newborn FGR (birthweight <10^th^ centile) and normally grown piglets were collected on day of birth and oral ibuprofen was administered daily for 3 days (20mg/kg/day, day 1 and 10mg/kg/day days 2 and 3). FGR brains demonstrated an inflammatory state, with changes to glial morphology (astrocytes and microglia), and blood brain barrier disruption, assessed by IgG and albumin leakage into the brain parenchyma and a decrease in blood vessel density. Loss of interaction between astrocytic end-feet and blood vessels was evident in the microvasculature where leakage was present, suggestive of structural deficits to the NVU. Ibuprofen treatment reduced the pro-inflammatory response in FGR piglets, reducing levels of pro-inflammatory cytokines and number of activated microglia and astrocytes associated with blood vessels. Ibuprofen treatment also attenuated albumin and IgG leakage. There were no alterations to angiogenesis, or blood vessel proliferation in treated-FGR piglets. These findings suggest postnatal administration of ibuprofen on day of birth modulates the inflammatory state, allowing for stronger interaction between vasculature and astrocytic end-feet to restore NVU integrity. These changes to the FGR brain microenvironment may be key to neuroprotection.

## Introduction

Fetal growth restriction (FGR) is commonly caused by chronic placental insufficiency whereby inadequate blood flow interrupts the supply of oxygen and nutrients to the fetus compromising fetal growth. The chronic hypoxic environment in which the FGR fetus develops can significantly affect the developing brain and these infants exhibit a range of adverse long-term neurological outcomes from schooling and behavioural issues to cerebral palsy [12, 35, 43, 51]. Clinical imaging studies demonstrate alterations in grey and white matter volume and structure in FGR infants [26, 71, 88], which are associated with developmental disabilities [26, 88]. Determining underlying mechanisms behind brain impairment in the FGR newborn will facilitate the rational choice or development of therapies to protect the vulnerable FGR brain.

The neurovascular unit (NVU) is a multicellular component of the central nervous system (CNS) that serves to separate the brain from the blood. The NVU is composed of vascular endothelial cells, glial cells (astrocytes and microglia), neurons, pericytes, and the basement membrane. The cells of the NVU share close and complex interactions that are crucial in maintaining blood-brain barrier (BBB) integrity and cerebral homeostasis, ensuring healthy brain development. The NVU plays a critical role in protecting the brain against the entry of toxic substances, which can have long-term pathological effects on the brain. Disruption to the NVU has been shown to play an essential role in development and progression of numerous CNS pathologies in the adult brain, such as Alzheimer’s disease (AD), multiple sclerosis (MS), stroke, and ischemia [83, 85]. When the NVU is disrupted, toxic substances, such as pro-inflammatory cytokines and immune cells, can infiltrate the brain, perpetuate neuroinflammation and injure neurons and white matter [8, 83]. Recently, much research has been undertaken to understand the role of the NVU in adult disease states, there has been little research on the involvement of the NVU in perinatal insults.

The developing brain is particularly vulnerable to changes in microenvironment and thus any disruption to the NVU may result in detrimental consequences on the newborn brain. Inflammation may be a key mediator contributing to NVU dysfunction in neuropathological conditions [56]. We have shown a robust neuroinflammatory response in the FGR newborn brain with increases in activated microglia and reactive astrocytes in the brain parenchyma [92, 94]. Microglia and astrocytes are critical for normal brain function and while in their resting morphological state, have a close interaction with blood vessels and are involved in BBB function, regulation, and stability [1, 8]. However, when microglia and astrocytes are activated in response to injury, they change morphology and function, which can lead to NVU disruption, as evidenced in adult neurological animal studies [1, 31, 38]. Further to the disruption of structural integrity, these cells no longer act to support the NVU, releasing large amounts of pro-inflammatory cytokines, such as tumor necrosis factor-α (TNFα) and interleukin-1β (IL-1β), which are detrimental to the brain microenvironment [38].

The neuroinflammatory response in the FGR newborn piglet brain comprises an increase in activated microglia, reactive astrocytes and pro-inflammatory cytokines (TNFα and IL-1β), and associated neuronal and white matter impairment [92, 94]. Inflammation and brain impairment persist days after birth [92, 94]. Recent sheep studies demonstrate inflammation may be associated with NVU disruption in newborn FGR lambs 24h after birth [15, 16] [16, 54]. Deficits in various components of the NVU may play a significant role in brain injury seen in these FGR animals. Targeting treatments to protect the NVU may reduce or even prevent the influx of toxic substances, such as inflammatory mediators and immune cells, into the brain and protect the vulnerable newborn brain. Ibuprofen is a non-steroidal anti-inflammatory drug currently used in the preterm newborn to treat patent ductus arteriosus [70]. Ibuprofen inhibits cyclooxygenase enzymes (COX1&2) which have critical functions on pro-inflammatory processes. We recently demonstrated ibuprofen treatment reduces inflammation and neuronal and white matter disruption in the FGR piglet brain [94]. Microglia and astrocytes in the FGR brain changed in morphology to a resting state following treatment with a concurrent decrease in pro-inflammatory cytokines. Ibuprofen acts on inflammation and modifies glial responses in the injured newborn brain, but whether ibuprofen can protect the NVU in the FGR newborn is yet to be determined.

In the current study, we hypothesized that the NVU is compromised in the neonatal FGR brain as a result of neuroinflammation. Administration of an anti-inflammatory agent, ibuprofen, improves NVU integrity by modulating juxtavascular glia. To undertake this study, we used our preclinical model of FGR, which in pigs arises due to placental insufficiency, the most common cause of FGR in the human population [9]. We observed presence of endogenous blood-borne proteins and immune cells in the FGR brain that were associated with alterations to structural components of the NVU and mediated by the brain’s inflammatory environment. Our findings indicate NVU alterations in the FGR neonate are associated with early inflammatory responses in the brain and potentiation of sustained brain injury.

## Materials and methods

### Animals and tissue preparation

Ethics approval for this study was granted by The University of Queensland (MED/UQCCR/132/16/RBWH & UQCCR/249/18) and the study was carried out in accordance with the National Health and Medical Research Council guidelines (Australia) and ARRIVE guidelines. The piglet is an appropriate animal to examine altered brain development arising from compromising perinatal events [11, 62, 74]. The piglet brain is gyrencephalic and has a similar grey to white matter ratio [20] as well as brain growth spurt in the perinatal period similar to the human [23]. FGR in the piglet is highly relevant to the clinical newborn situation as FGR occurs spontaneously in the pig by placental insufficiency[9] as occurs in the human FGR newborn.

Newborn Large White FGR (<10^th^ percentile birth weight; n=14) [41, 92] and normally grown (NG) piglets (n=14) (<18 h) were collected from the UQ Gatton Piggery on the day of birth, monitored and cared for at the Herston Medical Research Centre (HMRC) until day of euthanasia on postnatal day 4 (P4). Litter-matched piglets were obtained from multiple sows (n=12) with equal males and females in each group.

Animals were assigned to one of four groups: NG=8, FGR=8, FGR+ibuprofen=6, NG+ibuprofen=6. Ibuprofen treatment groups received an oral dose of 20mg/kg/day of ligquid ibuprofen on day 1 and 10mg/kg/day on days 2 and 3, as previously described [94]. This dosage is similar to that routinely used clinically in the human preterm newborn to treat patent ductus arteriosus [70], and is effective at reducing inflammation in the FGR piglet brain [94]. On P4, piglets were euthanased via an intraperitoneal injection of sodium phenobarbital (650mg/kg; Lethabarb, Virbac, Australia). The brain was transcardially perfused with physiological saline then collected, weighed, hemisected and coronally sliced. Tissue slices from the right hemisphere containing the parietal cortex were immersion fixed in 4% paraformaldehyde. Parietal cortex from the left hemisphere was snap frozen in liquid nitrogen and stored at −80°C for protein analysis as previously described [40].

### Magnetic Resonance Imaging

#### Preparation of animal

On postnatal day 3, piglets were sedated using Zolteil (Tiletamine/Zolazepam 10 mg/kg intramuscular injection, Virbac, NSW, Australia) and cannulated in the mammary vein (ease of access) using 50 cm of tygon tubing (ID 0.02 inch, OD 0.06 inch) with just the metal needle from a 25G needle on one end and an inserted 25G needle on the other end. Each piglet was then placed in a 300mm bore 7T ClinScan MR scanner (Bruker, Germany), running Siemens VB17. A 150 mm ID MRI rf coil was used to acquire the dynamic images. A syringe was prepared and connected to the cannula containing 1mL Gadovist 1.0 (Bayer) and connected to the cannula.

#### MR-PET Imaging procedure

The dual echo dynamic gradient echo (GRE) images were acquired with the following parameters: GRE_5slice: flip angle = 45°, 5 X 2 mm slices, FOV = 79 X 69 mm, in-plane resolution = 411 x 411 µm, TR = 50 ms, TE = 3 and 6 ms, acquisition time = 6.2 sec, 120 repetitions, total imaging time 12 min 33 sec. After a 2 min baseline period the Gadovist solution was injected.

#### Image analysis and calculations

HOROS (www.horosproject.org) was used for MRI image visualisation and region-of-interest (ROI) drawing and analysis. On the central slice, one ROI was positioned within the thalamus and one in the cortical regions (Figure 1) for the TE = 3 and TE = 6 image sets. The ROIs were then propagated to images from all time points, then exported as a csv files and imported into excel for calculations. Calculation of the gadolinium concentration derived independently from T2* and T1 time changes were determined from equations modified as previously described [46], using TE = 3 and TE = 6, r2* = 6.35 and r1 = 6.1 (Gadovist solutions at 7.0 T), A= mean ROI intensity (TE = 3) and B = mean ROI intensity (TE=6), M_o_sinα = 2454. T_2_* derived concentration = ((1/(3/ln(A/B)))-(1/(T_2_*_baseline_)))/6.35. T_1_ derived concentration = ((1/(abs(75/(ln((1-(A*(A/B)/2454))/(1-(A*(A/B)/2454 *0.707)))))) - (1/T_1 baseline_))/6.1). The values were then expressed as change from the baseline and a 3 point floating average was applied.

**Fig. 1.**
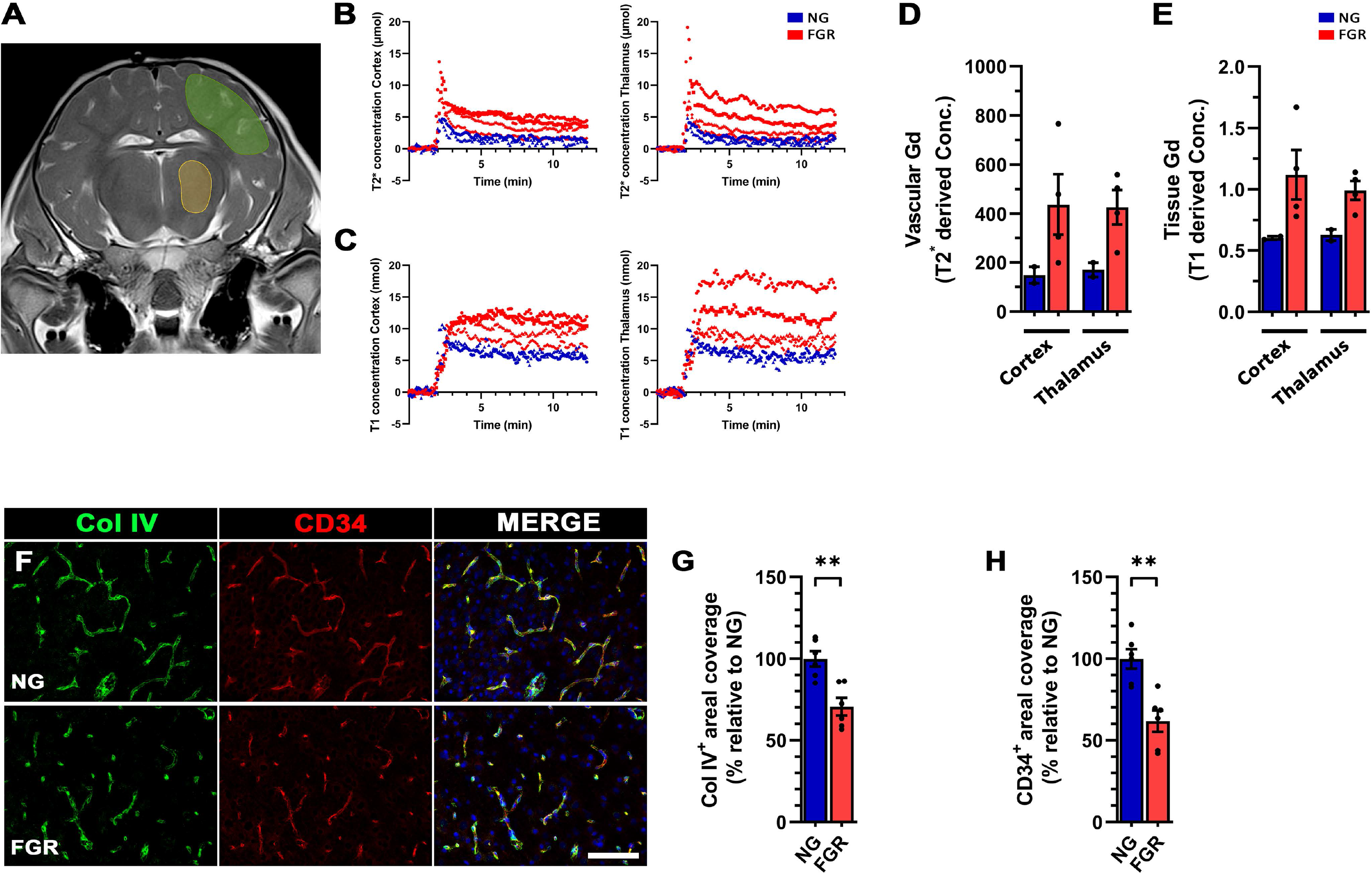
FGR neonates display altered vascular integrity and morphology. (***A***) Gadolinium-contrast MRI did not demonstrate overt leakage discernible by visual inspection in the FGR brain at postnatal day 3. Time course analysis demonstrated altered handling of gadolinium (Gd) in FGR brain (red) compared with NG (blue) over the 12min dynamic recording period. (***B***) T2* concentration in the cortex and thalamus displayed a rapid rise following administration followed by gradual decay. (***C***) T1 concentration demonstrated a maintained plateau that was evidently higher in FGR than observed in NG cortex and thalamus. Quantification of dynamic gradient echo (GRE) images found higher levels of Gd in the vasculature (***D***) as well as the tissue (***E***) in the thalamus and cortex of FGR. (***F***) Representative labelling of vascular basement membrane protein (green; Col IV) and endothelial cells (red). FGR brains displayed more truncated and discontinuous labelling patterns of Col IV and CD34 compared with NG. (***G*** *& **H***) Quantification of areal coverage confirmed a decrease in both Col IV and CD34 in FGR brains compared with NG. All values are expressed as mean +/− SEM (for vessel analysis; minimum *n* = 6 for NG and FGR). Unpaired student’s t-test (\*\**p* < 0.01) (Scale bar: 100μm).

### Immunohistochemistry

Fixed brain sections containing parietal cortex were embedded in paraffin and coronally sectioned in 6µm slices. For vessel structure and glial interaction analyses, brain slices of 12µm thickness were used. Sections were affixed to Menzel Superfrost Plus adhesive slides and dried overnight at 37°C. All sections were dewaxed and rehydrated using standard protocols followed by heat induced epitope retrieval with 10mM citrate buffer (pH 6) or TRIS-EDTA buffer (pH 9) at 90°C for 20 min before cooling to room temperature (RT). Tissue was blocked with 5% donkey serum in PBS with 0.5% Triton-X 100 for 1h at RT. Primary antibodies (Supplementary Table 1) were incubated overnight at 4°C. Slides were washed in tris-buffered saline followed by incubation with species-specific secondary fluorophores (Supplementary Table 1) at RT for 1h. Brain tissue was then washed, counterstained with 4′,6-diamidino-2-phenylindole (DAPI), and mounted with Prolong Gold antifade (Molecular Probes, Invitrogen Australia, Victoria, Australia). Negative control sections without primary antibodies were processed in parallel and immunolabelling was conducted in triplicates for all animals.

For angiopoietin labelling, a tyramide signal amplification (TSA) kit was used following the manufacturer’s instructions (Perkin Elmer; NEL791001KT). Briefly, slides were prepared as described above and incubated with either angiopoietin 1 (Ang1) or angiopoietin 2 (Ang2) overnight at 4°C. Slides were then washed, and endogenous peroxidase activity was quenched with 3% H_2_O_2_ for 5 minutes. Conjugated HRP secondary (1:2000) was incubated for 1h at RT, followed by addition of TSA-Cy5 complex. Slides were washed and the alternate Ang antibody added to incubate overnight at 4°C. After washing, slides were incubated with Alexafluor-488 and DAPI for 1h at RT, washed and mounted as described above.

### Image acquisition and analysis

Analysis of immunolabelled sections was performed using a Zeiss Axio Microscope with an Axiocam503 camera. Four pictomicrographs of the parietal cortex were captured for analysis from each slice. For all markers, replicates were conducted with slices separated by at least 100μm. All imaging and analyses were conducted under blinded conditions by KKC and JAW. Microglia were manually counted and categorised with respect to morphology as per previous studies [92, 94]. Density analysis, co-localisation and vessel coverage analyses was undertaken using the threshold function with moments plugin in FIJI (ImageJ; Image Processing and Analysis in Java; National Institutes of Health, Bethesda, MD, USA).

### Western blotting

For quantification analyses of tight junction proteins (claudin5, occludin, zonula occludin) [28] and angiogenesis (Ang1 and Ang2), brain tissue from the parietal cortex was homogenised in 5× volume of 1M Tris-HCl-EDTA (0.5M) buffer and centrifuged at 6,300 X *g* for 10min at 4°C. The supernatant was collected, and bicinchoninic acid (BCA) assays were performed to estimate total protein concentrations (Life Technologies; #23225). Total protein lysates (5-20 µg) were resolved on either 7%, 10% or 12.5% sodium dodecyl sulphate polyacrylamide (SDS-PAGE) gels at 150V for 90min; dependent on predicted molecular weight for protein. Proteins were transferred to a PVDF membrane (0.45µm Immobilon^®^-P, Merck Millipore Ltd., VIC, Australia) at 100 V in chilled transfer buffer containing 25 mM Tris, 40 mM glycine and 15% methanol for 70min at 4°C. The membranes were blocked with 5% blocking buffer (non-fat skim milk powder) prepared in Tris-buffered saline with 0.1% Tween-20 (TBST) at RT for 1h. Membranes were incubated overnight with either anti-claudin-5 (Cldn5; 1:100,000), anti-occludin (OCLN; 1:6,000), anti-zonula occludin (ZO1; 1:7,500), anti-Ang1 (1:10,000), or anti-Ang2 (1:10,000). Membranes were washed and incubated in horseradish peroxidise-conjugated anti-rabbit IgG secondary antibody (1:50,000 OCLN and ZO1; 1:80,000 Cldn5, Ang1 and Ang2) for 1h at RT. Protein signal was developed using enhanced chemiluminescence according to the manufacturer’s instruction (Immobilon^®^ Forte Western HRP substrate, Merck Millipore Ltd., VIC, Australia) and visualised using the ChemiDoc^TM^ XRS system (Bio-Rad; Hercules, USA). Each membrane was washed twice with 0.1% TBST and re-probed with anti-β-actin (1:50,000; ab8225) to verify uniform quantity of protein loaded on each blot. Each antibody was quantified relative to β-actin levels as previously described [93] using ImageJ software (National Institutes of Health, MD, USA).

### Statistics

All statistical analyses were performed using Graph Pad Prism 9.0 software, San Diego, California, USA. Data were analyzed using either Unpaired Student’s t-test or a two-way ANOVA with Tukey post-hoc analysis. Results were expressed as mean ± SEM with statistical significance accepted at p<0.05.

## Results

At postnatal day 4, (P4) mean body weight was significantly lower for both untreated FGR (p<0.0001; p<0.01) and treated FGR piglets (p < 0.0001; p<0.01) compared with untreated NG and treated NG piglets respectively (Table 1). Brain weight was significantly reduced in both the untreated-FGR piglets (p = 0.009) and FGR-treated piglets (p = 0.031) in comparison to untreated NG piglets. However no significant difference in brain weight was evident in untreated-FGR (p = 0.061) and treated FGR (p = 0.148) piglets compared to treated NG piglets. Asymmetric FGR piglets were defined by birth bodyweight (<10^th^ percentile on the day of birth) and by brain to liver weight ratio (B:L) ≥ 1 on P4 [10, 21, 41]. There was no significant difference in body weight (p = 0.234), brain weight (p = 0.996) or liver weight (p = 0.780) between treated and untreated NG piglets (Table 1).

**Table 1:** Piglet bodyweight, brain weight and liver weight

|  | **NG (n=8)** | **FGR (n=8)** | **FGR+ibuprofen (n=6)** | **NG+ibuprofen (n=6)** |
| --- | --- | --- | --- | --- |
| Bodyweight in grams<br>(mean±SEM) | 1968 ± 138.10 | ††1018 ± 74.68**** | ††1000 ± 39.67**** | 1643 ± 125.6 |
| Brain weight in grams<br>(mean±SEM) | 32.74 ± 0.50 | 29.97 ± 0.68** | 30.18 ± 0.49* | 32.29 ± 0.59 |
| Liver weight in grams<br>(mean±SEM) | 51.24 ± 6.16 | 25.92 ± 2.80** | †22.69 ± 2.09*** | 43.44 ± 4.76 |
| Brain:liver ratio<br>(mean±SEM) | 0.69 ± 0.07 | †1.249 ± 0.14** | †1.383 ± 0.12** | 0.797 ± 0.10 |

Piglet bodyweight, brain weights and liver weight. Mean body weight was significantly lower in both FGR untreated and ibuprofen treated piglets compared to untreated NG piglets. Brain weight was significantly reduced only in the FGR untreated and treated groups compared to the NG group. The mean brain to liver weight ratio was significantly higher in both treated and untreated FGR piglets compared to NG piglets indicating asymmetric growth restriction in the FGR piglets. Values are the mean±SEM. \*p<0.05; \*\*p<0.01; \*\*\*p<0.001; \*\*\*\*p<0.0001 versus NG. †p<0.05; ††p<0.01 versus NG+ibuprofen.

### Perturbed vascular integrity in FGR newborn brains

Magnetic resonance imaging (MRI) with gadolinium-based contrast agents has been used for many years in patients with a spectrum of CNS disorders associated with NVU disruption such as stroke and multiple sclerosis. Here, we utilised MRI with gadolinium (Gd) to examine whether there was a detectable *in vivo* alteration to NVU function in the cortex and thalamus of FGR newborns at postnatal day 3 (Fig. 1). The T2*-derived Gadovist concentration, estimates high Gd concentrations (µmol) (Fig. 1B). This is typically reflected by the rapid increase of vascular Gd due to the bolus passing through the tissue and then equilibration of the Gd throughout the total blood volume. The T1-derived Gadovist concentration, estimates lower Gd concentrations (nmol) (Fig. 1C), displays a slower uptake, followed by a plateau with gradual decrease, consistent with Gadovist crossing the BBB and entering the brain tissue [34]. The total vascular and tissue Gd was estimated over the 12 minutes of dynamic acquisition. Total vascular Gd was estimated from the summation of all data points from T2* derived concentration curves while total tissue Gd was estimated from T1 derived curves. We observed altered Gd uptake and clearance rates in both regions investigated in the FGR brain compared with NG following Gd injection (Fig. 1B & C). FGR displayed an elevated concentration of vascular and tissue Gd in both the cortex and thalamus (Fig. 1D & E).

Given that these findings suggest alterations in vascular function, we next examined gross vessel structure using collagen IV (Col IV), which makes up approximately 50% of the basement membrane [50], and a marker of endothelial progenitor cells (CD34) (Fig. 1F). FGR displayed positive labelling for both markers, however a significant decrease in both Col IV (*p =* 0.013) and CD34 areal density (*p* = 0.001) was found suggestive of lower vessel density in FGR compared with NG (Fig. 1G & H).

Investigation of proliferative cells (Ki67-positive) showed a lower number in both the parenchyma (*p* = 0.028) and juxtavascular (*p =* 0.002) regions of FGR brain, with further analysis showing a decrease in the number of Ki67-positive vessels (Supp. Fig. 1A-C). There was no overt difference in labelling for angiopoietin 1 and 2, markers of angiogenesis (Supp. Fig. 1D). Western blot analysis found no significant difference between FGR and NG for either Ang1 (*p* = 0.140) or Ang2 (*p* = 0.305) (Supp. Fig 1E & F). Analysis of Ang2 to Ang1 ratio, which may indicate a pro-angiogenic state, showed no significant difference between FGR (1.42 ± 0.07) and NG (1.18 ± 0.11) (*p* = 0.082) (Supp. Fig. 1G). Together these findings suggest altered vascular integrity which may be associated with perturbed NVU composition.

### Pro-inflammatory glia interact with vasculature of FGR newborn brains

We have previously reported postnatal neuroinflammation associated with reactive glial morphology in the FGR brain [92, 94]. Here, we specifically sought to understand if glial cells that interact with the brain vasculature demonstrated altered morphology in the FGR brain. In NG animals, both astrocytes and microglia near blood vessels (juxtavascular) demonstrated a resting morphology, with long thin process extensions and rounded cell bodies (Fig. 2B-b’ & Supp. Fig. 2A) while astrocyte end-feet interacted closely with the vasculature and spanned the majority of the microvessels (see Fig. 2B for cross-sectional interaction & Supp. Fig 2A for transverse representation). In the FGR brain there was increased juxtavascular glial activation (Fig. 2C; Supp. Fig. 2B), as well as an increase in the number of Iba-1 positive microglia (Fig. 2G). There was evidence of altered astrocyte end-feet contact with the vessel and in regions of the brain with severe injury, there was overt loss of astrocyte end-feet interaction with the vessel in FGR brains (Fig. 2D & Supp. Fig. 2C).

**Fig. 2.**
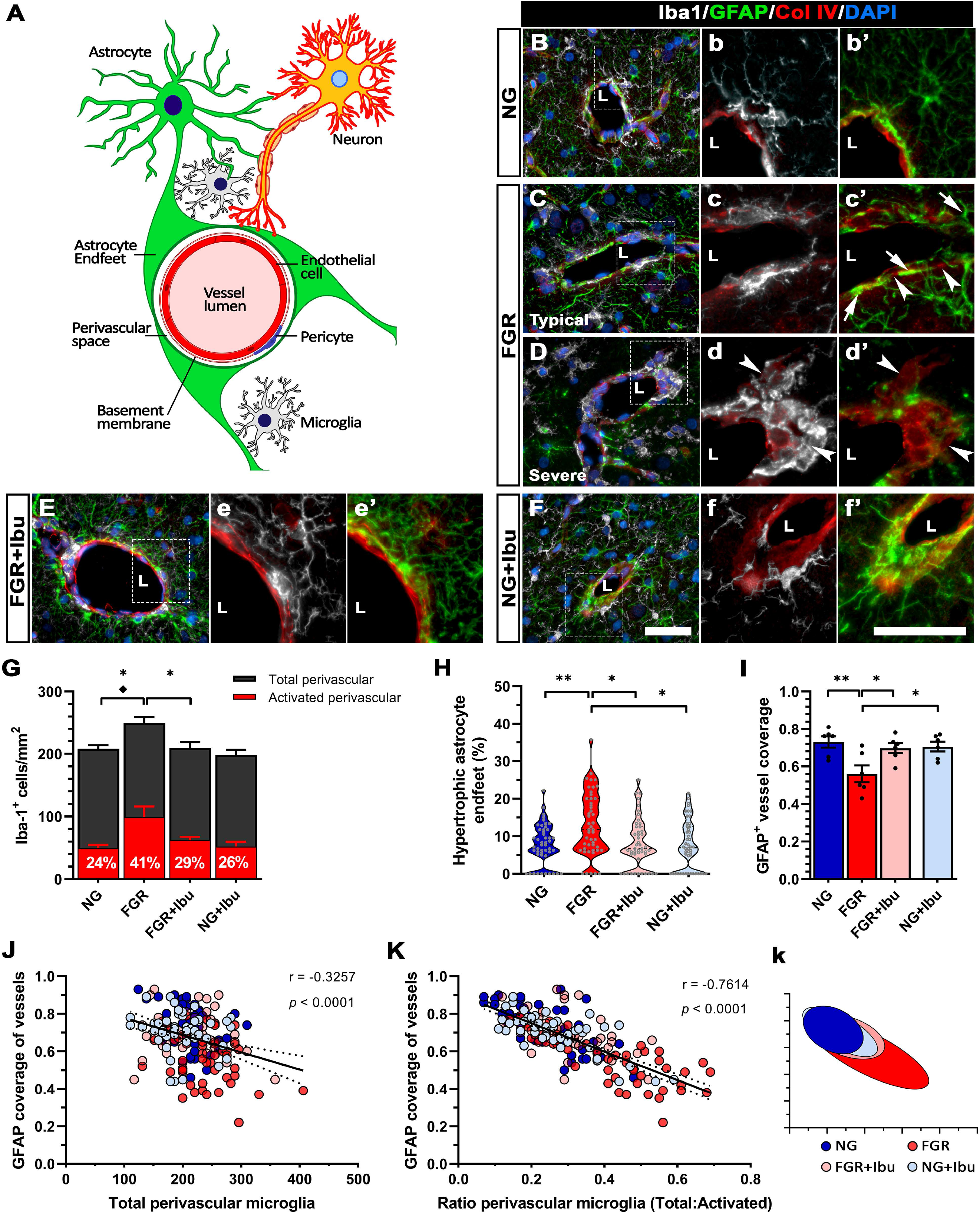
Activated glia interact with microvessels of FGR newborns. (***A***) Schematic illustrating the cellular composition of the healthy neurovascular unit (NVU). Neuronal and glial cells signal to maintain NVU integrity and provide a homeostatic brain environment. (***B***) Representative labelling of microglia (grey) and astrocyte (green) interactions with a vessel (lumen; L) in the NG brain in a cross-sectional plane. NG demonstrated resting glial morphology, with long thin process extensions forming an organised network of consistent interactions with the vasculature (see ***b*** & ***b’***). (***C***) FGR brain showed activation of glia based on morphology, with retraction of extended processes and dense cellular bodies. Cross-sectional micrograph demonstrates close interaction of activated microglia with the vessel (***c***), high intensity stained astrocyte end-feet (***c’*** arrows), and regions sparse of GFAP labelling along the vessel margin (***c’*** arrowhead). (***D***) Instances of severe alterations in glia interactions were noted in FGR brain, with dense clustering of activated juxtavascular microglia (***d***) ***s***ignificant loss of GFAP labelling around vasculature (arrowheads; ***d’***). (***E*** & ***F***) Ibuprofen treated FGR and NG displayed morphology comparable to untreated NG brains, indicating a return to resting state for glial cells. (***G***) Quantification of labelling demonstrated an increase in total microglia as well as an increase in activated juxtavascular microglia in FGR brain relative to NG. (***H***) Analysis of GFAP labelling found an increase in hypertrophic end-feet and (***I***) a decrease in GFAP^+^ vessel coverage in the FGR brain which was ameliorated by ibuprofen treatment. (***J***) Correlative analysis found no significant relationship between total microglia and GFAP^+^ vessel coverage. (***K***) Analysis showed an inverse correlation between GFAP^+^ vessel coverage and the ratio of total to activated juxtavascular microglia, with FGR skewed to the right (***k***). All values are expressed as mean +/− SEM (minimum *n* = 6 for all groups). Two-way ANOVA with Tukey post-hoc test (\**p* < 0.05; \*\**p* < 0.01) (Scale bars: 50μm).

In ibuprofen treated FGR animals, juxtavascular astrocytes and microglia displayed resting morphology similar to that observed in brains of NG animals (Fig. 2B,E,F & Supp. Fig. 2A,D,E). FGR brains displayed an increased frequency of hypertrophic astrocytic end-feet (determined by enlarged, dense, high-intensity labelling at the vessel interface - Fig. 2c’,d’) compared with NG animals (*p* = 0.003), which corresponded to a decrease in astrocytic endfeet coverage of blood vessels (Fig. 2D&I: *p* = 0.007). Ibuprofen treatment ameliorated these changes reducing frequency of hypertrophic astrocytic end-feet (Fig 2H; *p* = 0.046) and normalising vessel coverage (Fig. 2I; *p* = 0.036). Treatment with ibuprofen also reduced the number of activated juxtavascular microglia in FGR animals (Fig 2G; *p* = 0.033). Correlative analyses demonstrated a negative relationship between astrocyte vessel coverage and the number of perivascular microglia (r = −0.3257) (Fig. 2J). We also found a negative correlation between astrocytic vessel coverage and the number of activated perivascular microglia (r = − 0.7342) (Fig. 2K & k). These findings suggest that activated microglia influence the interaction of astrocytes with the brain vasculature in the newborn.

Activation of microglia is associated with increased expression of pro-inflammatory numerous cytokines (Fig. 3A) Our previous findings demonstrated elevated levels of several pro-inflammatory cytokines in the FGR brain parenchyma which was ameliorated following three days of ibuprofen treatment [94]. In the current study, we specifically examined the expression of these inflammatory mediators at the NVU. FGR animals showed robust labelling of NF-κΒ, TNFα, and IL-1β in microglia and astrocytes as well as co-located neurons compared with NG animals (Fig. 3B, C & D). Treatment with ibuprofen reduced staining for these markers while there was strong labelling for the anti-inflammatory mediator IL-4 (Fig. 4A). Labelling for CXCL10 (IP10), a known mediator of BBB-disruption, showed strong expression at the vasculature of FGR animals as well as in microglial cells and neurons (Fig. 4B). CCL2 was robustly expressed in neuronal cells (Fig. 4C), while CCL3 stained microglia (Fig. 4D), which appear to engulf CCL3^+^ neurons in the FGR brain (Fig.2H). The expression of CCL2 and CCL3 was sporadic and restricted in NG and ibuprofen treated groups.

**Fig. 3.**
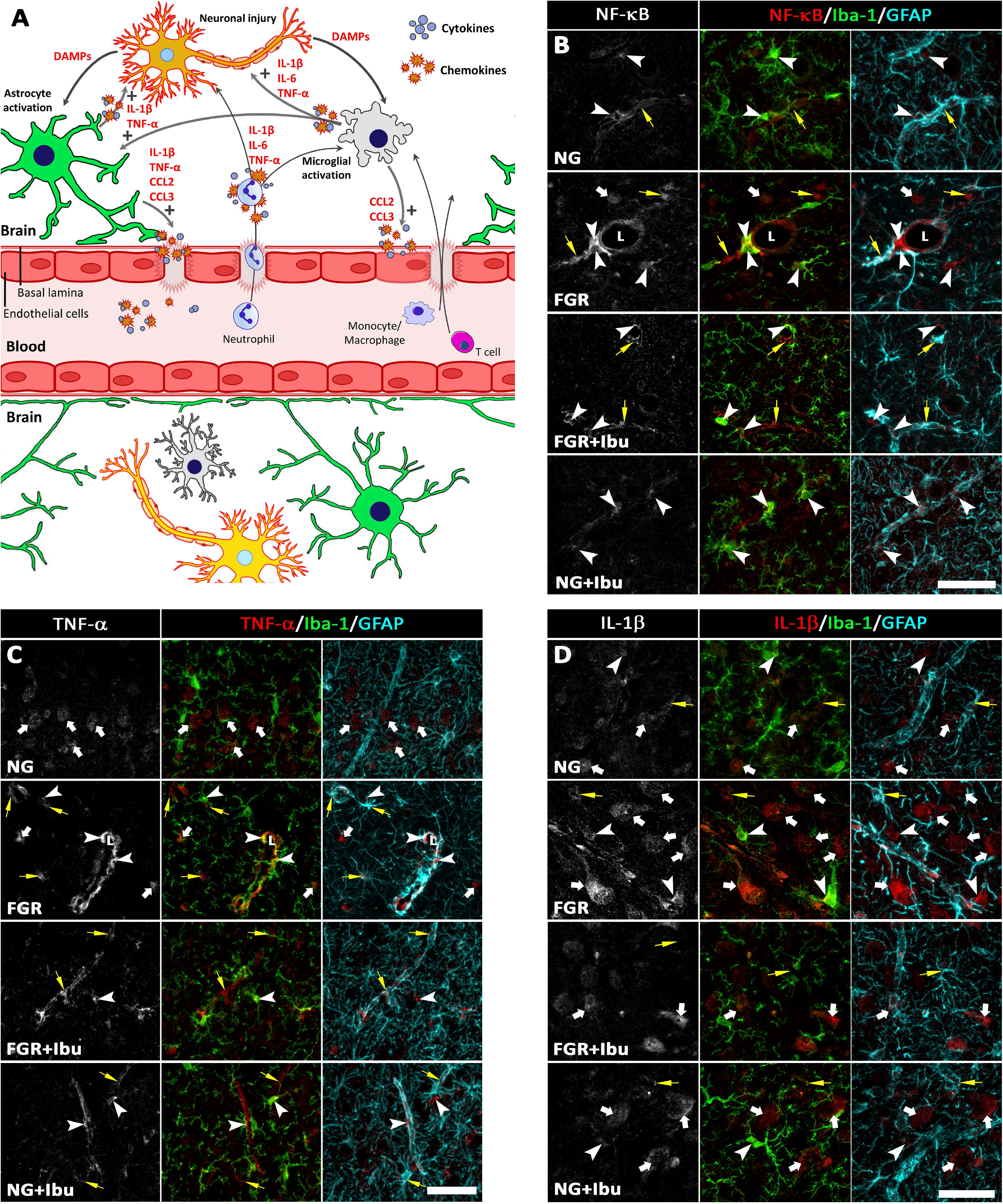
Increased expression of pro-inflammatory mediators at the vasculature of FGR brains. (***A***) Schematic demonstrating the interplay between glial cell activation and the release of cytokines and chemokines which can further NVU-disruption and promote the adhesion of circulating peripheral immune cells. Infiltration of these cells through the permeable NVU can further exacerbate the already established pro-inflammatory environment of the FGR newborn brain. Representative labelling of pro-inflammatory mediators which were robustly expressed in cells comprising the NVU in FGR brain. Classical pro-inflammatory mediators (***B***) NF-κΒ, (C) TNFα, and (***D***) IL-1β demonstrated strong co-localisation with microglia (Iba-1; arrowheads), astrocytes (GFAP; yellow arrows), neurons (white arrows) and vasculature. NG, FGR+Ibu and NG+Ibu brains showed sporadic, low intensity labelling for each pro-inflammatory mediator investigated in comparison with FGR littermates. A minimum of *n* = 5 animals were stained for each marker per group (Scale bars: 50μm).

**Fig. 4.**
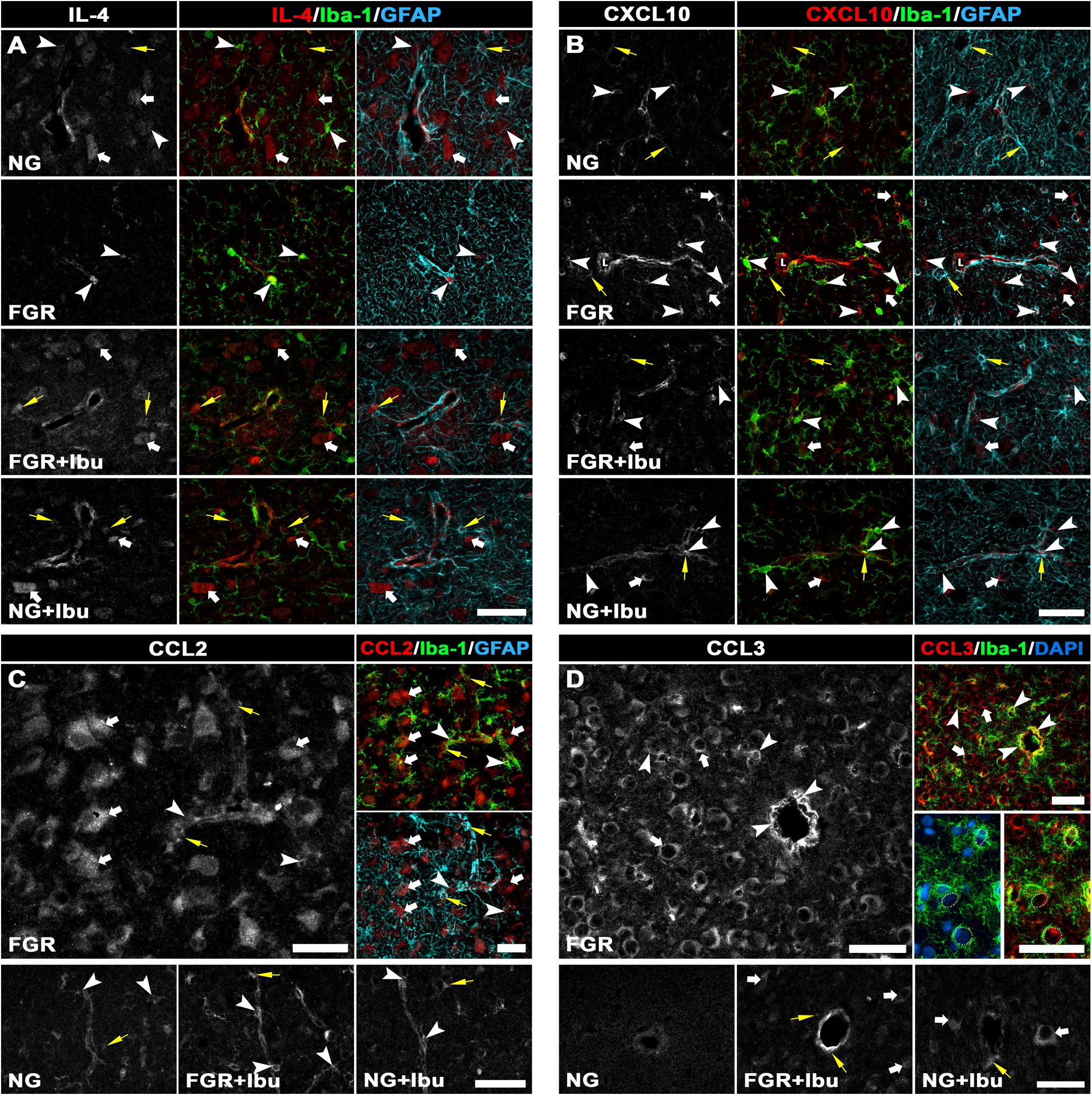
Expression of inflammatory mediators in glial, neuronal and vascular cells at postnatal day 4. (***A***) NG and ibuprofen treated groups demonstrated robust labelling of the anti-inflammatory cytokine IL-4 in neuronal cells (white arrows) and moderate expression in glia (microglia; arrowheads & astrocytes; yellow arrows). FGR brain showed low intensity labelling for IL-4 in microglial cells. (***B***) CXCL10 (IP10) was elevated in FGR compared with all other groups. The potent chemo-attractants for monocyte and immune cell recruitment, CCL2 (***C***) and CCL3 (***D***), were robustly expressed in neuronal cells and activated microglia in FGR brain. NG, FGR+Ibu and NG+Ibu brains showed sporadic, and relatively low intensity labelling for both CCL2 and CCL3. A minimum of *n* = 5 animals were stained for each marker per group (Scale bars: 50μm).

### Juxtavascular glia mediate entry of endogenous plasma proteins in FGR brain

The endogenous proteins albumin and IgG are useful in detecting BBB-disruption. We first examined the expression of the large endogenous protein IgG (∼155 kDa) which showed sporadic incidences of extravasation into the brain parenchyma with limited perivascular labelling (Fig. 5A & B, see E for surface plot analysis). In FGR animals IgG extravasation was observed in both the cortex (Fig. 5A) and underlying white matter (Fig.5B). Extravasation of IgG was observed at vessels with altered astrocyte interaction (loss of GFAP-positive labelling; see Fig. 5a) and in some instances, uptake of IgG by astrocytes appeared to preserve vessel interaction (Fig. 5b). There was only low intensity focal extravasion of IgG in FGR+Ibu and NG animals with astrocytes appearing to cordon off the leak (Fig. 5C & D respectively). Quantification of FGR and FGR+Ibu animals, showed ibuprofen administration ameliorated the increased IgG extravasation (*p* = 0.003; Fig. 5F), IgG areal coverage (*p* < 0.0001; Fig. 5G), and the intensity of IgG labelling relative to untreated FGR (*p* = 0.002; Fig. 5H).

**Fig. 5.**
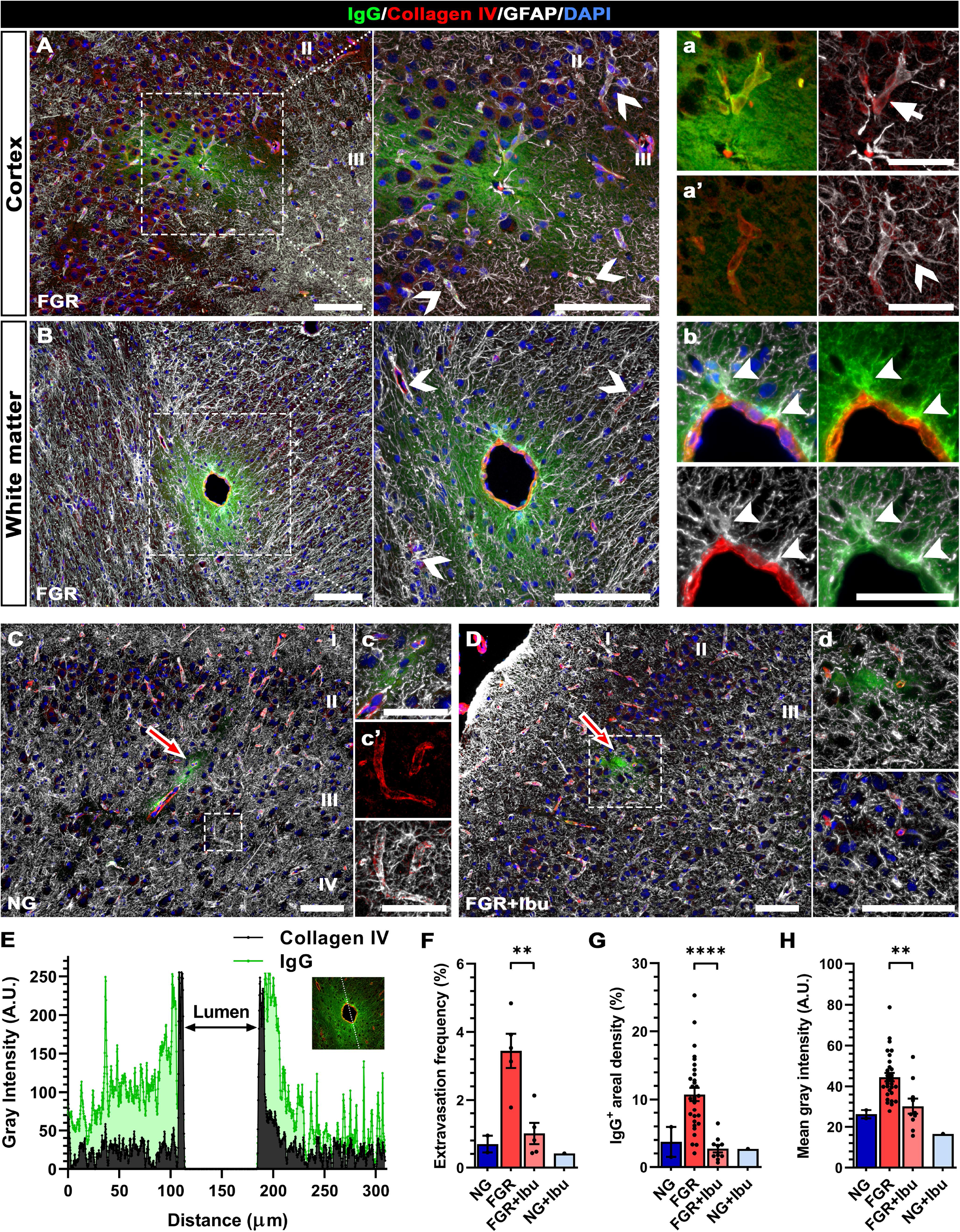
Ibuprofen treatment reduces frequency of IgG extravasation in FGR brain. Representative labelling of IgG (green), Collagen IV (vessels; red) and astrocytes (GFAP; white) expression. (***A***) FGR show IgG extravasation into the parenchyma of the cortex. Evident decrease in GFAP-positive labelling around vessels with extravasation (***a***; see arrow), while vessels with strong GFAP expression showed no IgG extravasation (***a’***; see chevrons). (***B***) IgG extravasation was also observed in white matter of FGR brains, with IgG uptake in astrocytes interacting with vessels at the glia limitans (***b***; see arrowheads). Both NG (***C***) and FGR+Ibu (*D*) showed infrequent focal IgG extravasation in the cortex and consistently displayed strong interaction of astrocyte end-feet around vessels and focal leaks (***c****, **c’*** & ***d***). (***E***) Example surface plot of IgG extravasation from the lumen of a vessel in FGR animals. Ibuprofen administration decreased the frequency of IgG extravasation (***F***), IgG areal coverage (***G***), and mean intensity of IgG labelling (***H***) when compared with untreated FGR. All values are expressed as mean +/− SEM. NG and NG+Ibu were excluded from analysis as limited animals showed presence of IgG, 2 and 1 respectively. For ***G*** & ***H***, each dot represents individual observations. FGR (*n* = 6) vs FGR+Ibu (*n* = 5) unpaired Student’s t-test, ** *p* < 0.01; **** *p* < 0.0001. (Scale bars: ***A***, ***B***, ***C*** & ***D-d***: 100μm; ***a-a’***, ***b*** & ***c-c’***: 50μm).

We observed three distinct patterns of albumin (∼69 kDa) labelling in the FGR brain. Perivascular labelling, where albumin was observed in the space between the basement membrane and astrocytic endfeet (Fig. 6A & B), uptake into juxtavascular astrocytes surrounding the vessels and sporadic extravasation with diffuse but decreasing labelling distal to the vessel (Fig. 6A & C). Albumin extravasation was associated with loss of astrocyte end-feet contact with the NVU (Fig. 6H, see chevrons). Where albumin was observed in juxtavascular astrocytes (Fig. 6h & h’; see arrows) or in the perivascular space (Fig. 6B,b & C,c), astrocyte endfeet maintained contact with the vessel but presented with hypertrophic end-feet indicative of a reactive morphology (Fig. 6h & h’; see arrowheads). Quantification of perivascular albumin labelling was evident on a greater number of vessels in FGR animals compared with NG animals (Fig. 6G; *p* < 0.0001). Treatment with ibuprofen reduced albumin extravasation (Fig. 6D-F) and significantly reduced perivascular labelling in FGR animals compared with untreated FGR animals (Fig. 6G; *p* = 0.0008), with the protein predominantly confined to the lumen of the vessel, and even distribution of astrocytic end-feet across the vasculature. These findings suggest that altered astrocyte interactions at the NVU contribute to altered BBB-permeability in the FGR brain (Fig. 6I).

**Fig. 6.**
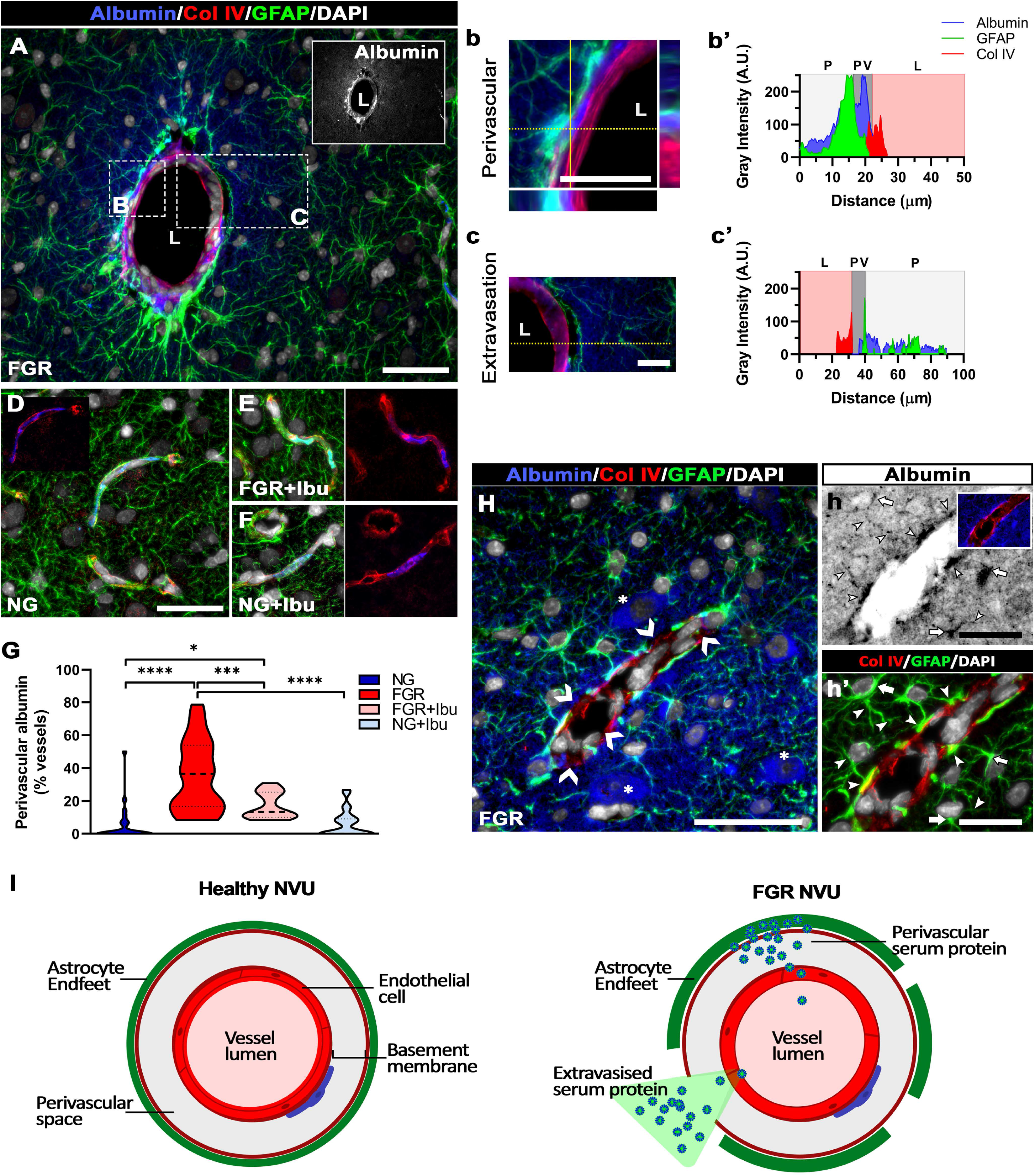
Increased perivascular albumin labelling in FGR brain. (***A***) Labelling of albumin (blue), basement membrane marker collagen IV (Col IV; red), and astrocytes (GFAP; green) in the FGR brain. (***B-b’***) Albumin was predominantly labelled in the perivascular space, between the basement membrane and astrocyte end-feet, with sporadic instances of (***C-c’***) extravasation into the brain parenchyma. (***b’***) Surface plot analysis demonstrating the high signal intensity of albumin labelling in the perivascular space (PV) located between the lumen (L) and parenchyma (P), while sites of extravasation demonstrated low intensity leakage tens of microns from the lumen (***c’***). (***D-F***) NG and treated groups demonstrated instances of albumin labelling restricted to the lumen of the microvessels. (***G***) FGR showed a significantly higher percentage of vessels with perivascular labelling compared with all other groups. (***H***) In FGR brains, regions with albumin extravasation and neuronal uptake (asterisk) also displayed decreased astrocyte end-feet interaction with vasculature, with some regions void of GFAP-labelling (***H***; see chevrons). (***h* & *h’***) Albumin uptake was observed in juxtavascular astrocyte end-feet and processes (arrowheads) as well as astrocyte bodies (arrows). (***I***) Schematic comparing healthy NVU composition with the proposed disrupted NVU in FGR. Astrocytic activation, hypertrophic end-feet and reduced vessel interaction at FGR NVU were associated with increased presence of endogenous serum proteins in the perivascular space, within end-feet and extravasation into the parenchyma. All values are expressed as mean +/− SEM (minimum *n* = 6 for all groups). Two-way ANOVA with Tukey post-hoc test (\**p* < 0.05; \*\*\**p* < 0.001; **** *p* < 0.0001) (Scale bars: ***A***: 100μm; ***D***-***F***, ***H***: 50μm; ***h*** & ***h’***: 25μm).

### BBB disruption results in T-cell infiltration in FGR newborn brains

Pro-inflammatory cytokines and chemokines released by cells of the NVU can enhance transmigration of peripheral immune cells, which in turn moderate inflammation and progression of disease pathology [82]. Using the pan T-cell marker (CD3), we examined T-cell infiltration into the FGR brain. CD3^+^ cells were present in the vessel lumen, as well as perivascular and the parenchyma of FGR brains (Fig. 7A). In comparison, NG and ibuprofen treated animals showed low CD3^+^ cell labelling which appeared to be localised to the microvessel lumen (Fig. 7B-D). Significantly elevated CD3^+^ cell counts were present in FGR when compared with NG (*p* = 0.041) and FGR+Ibu (*p* = 0.030) (Fig. 7E). Localisation analysis showed CD3^+^ in all groups were predominantly found in the vessel lumen, however FGR did display a shift toward increased perivascular and parenchymal localisation (Fig. 7F).

**Fig. 7.**
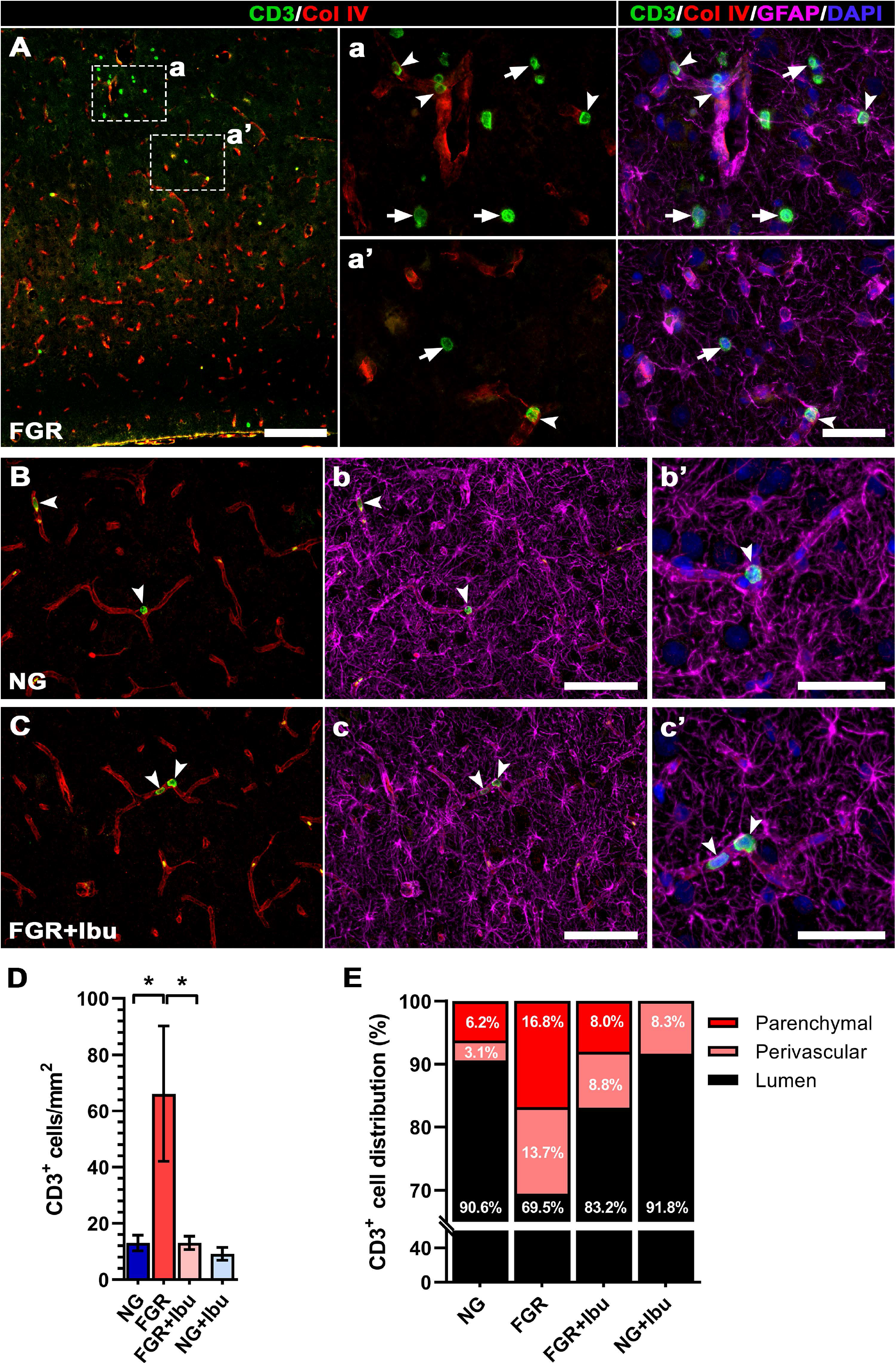
Altered NVU integrity results in immune cell infiltration into FGR brain. (***A***) Representative labelling of the pan T-cell marker (CD3) in FGR brain showed overt infiltration into the brain parenchyma (arrows), lumen and perivascular regions (***a*** & ***a’***; arrowheads). CD3^+^ cell infiltration was observed in regions presenting perturbed astrocyte-vessel interactions, as shown by altered GFAP labelling (magenta) around the vasculature (red). (***B***) NG and (***C***) FGR+Ibu demonstrated significantly lower numbers of CD3^+^ cells which were confined to the vessel lumen, with strong astrocyte-vessel interaction. (***D***) Quantification of immunofluorescent labelling found an almost three-fold increase in CD3^+^ cells in FGR compared with all groups. (***E***) A significantly higher proportion of these CD3^+^ cells were localized to the parenchyma and perivascular regions in FGR compared with NG. All values are expressed as mean +/− SEM (minimum *n* = 6 for all groups). Two-way ANOVA with Tukey post-hoc test (\**p* < 0.05) (Scale bars: ***A***: 200μm; ***B-b*** & ***C-c***: 100μm; ***a-a’***, ***b’*** & ***c’***: 50μm).

Infiltration of CD3^+^ cells is associated with increased expression of claudin-1 [4]. Claudin-1 expression increases in astrocytes which are thought to restrict CD3^+^ cells that have entered the brain parenchyma [4]. FGR displayed high expression of Cldn1 labelling in all layers of the cortex including the glial limitans at the pial surface (Supp. Fig. 3A). This was associated with the presence of CD3^+^ cells in the pial surface (Supp. Fig. 3Aa; see arrows), parenchyma (Supp. Fig. 3Aa & b), and perivascular space (Supp. Fig.3Aa & c). NG and FGR+Ibu displayed relatively low Cldn1 expression and limited CD3^+^ cell labelling (Supp. Fig. 3B & C). Analysis of Cldn1 expression showed elevated areal density in FGR compared with NG (Supp. Fig. 3D: p = 0.001). Astrocytic expression of Cldn1 was elevated in FGR brains (52.4%) compared with NG (13.9%; *P* < 0.0001) (Supp. Fig. 3E). FGR+Ibu treated animals displayed a 28.6% reduction in Cldn1 expression in astrocytes compared with untreated FGR (*p* = 0.001).

### Increased apoptosis in FGR newborn brain is ameliorated following ibuprofen treatment

We next investigated whether the altered BBB permeability in FGR brain was associated with apoptosis of cellular components of the NVU. FGR brain displayed elevated cleaved-caspase 3 (Casp3) labelling compared with all groups (*p* <0.0001; Fig. 8A&D), with approximately three times as many vessels displaying Casp3 labelling compared with NG (*p* <0.0001; Fig. 8D). Ibuprofen administration significantly decreased Casp3^+^ cells in FGR brain compared with untreated FGR (*p* <0.0001; Fig. 8A&D). FGR brain presented a higher number of Casp3^+^ cells per vessel compared with NG (*p* <0.0001; Fig. 8B&E), which was ameliorated following ibuprofen treatment (*p* <0.0001; Fig. 8E).

**Fig. 8.**
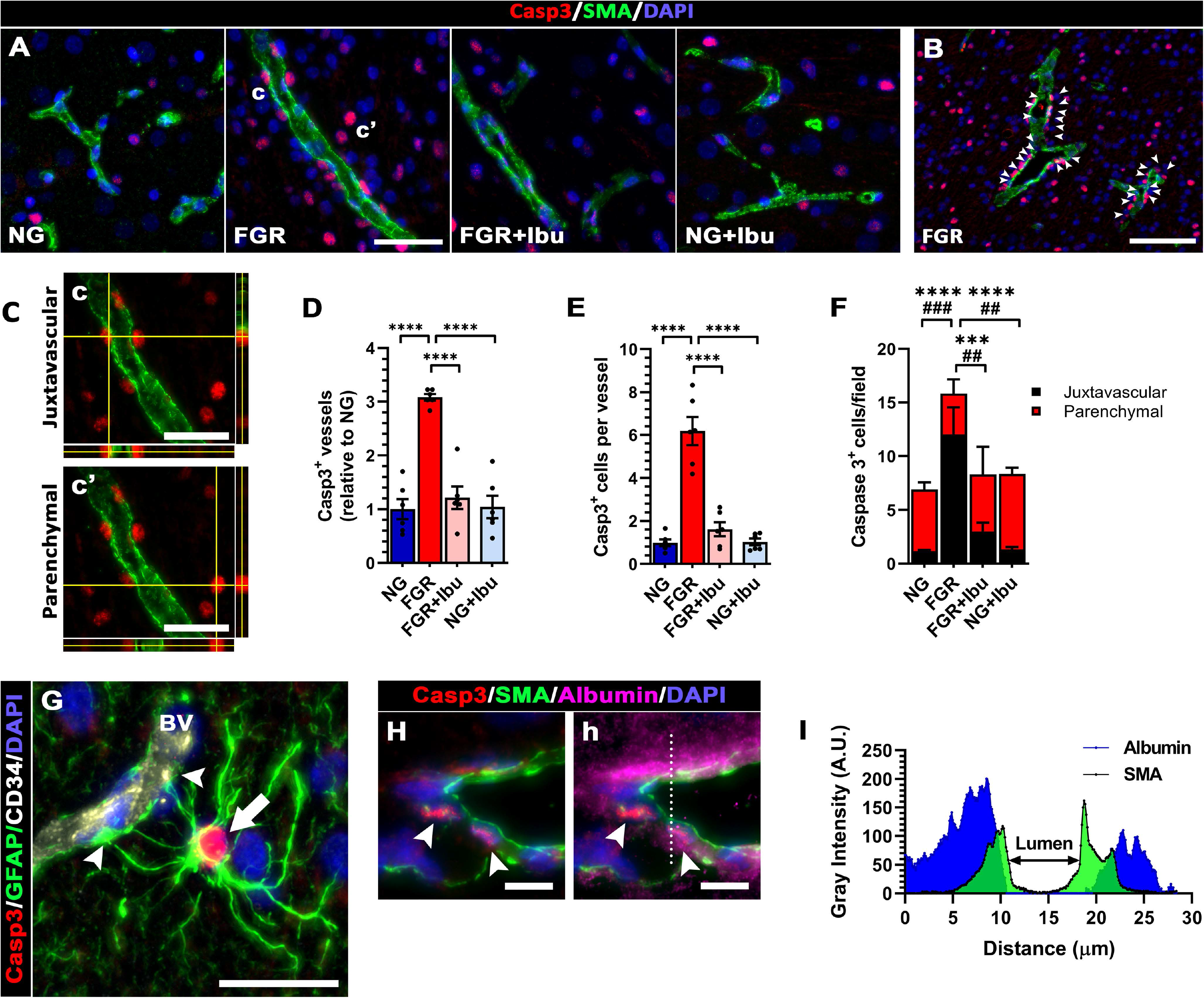
Ibuprofen treatment reduces apoptosis of cells comprising the NVU. **(*A*)** Representative labelling of apoptotic cells (Caspase 3; red) and vasculature (SMA; green) in NG, FGR, FGR+Ibu and NG+Ibu brain. (***B***) FGR brains displayed multiple Casp3-positive cells at individual vessels, as indicated by arrowheads. (***C***) FGR showed both juxtavascular (c; see close association between SMA and Casp3 in orthogonal views) and parenchymal (c’) caspase-3 labelling. (***D***) A threefold increase in the number of vessels expressing Casp3 was observed in FGR brains relative to NG. Ibuprofen treatment reduced the number of vessels with Casp3 expression. (***E***) FGR vessels had a higher number of Casp3-postivie cells per vessel when compared with all groups. (***F***) Increased juxtavascular and parenchymal Casp3 labelling was reported in FGR compared with NG. Ibuprofen treatment significantly reduced the number of Casp3 cells interacting with the NVU as well as in close proximity to the vessels. (***G***) Parenchymal Casp3 was strongly co-localised with astrocytes (GFAP) which displayed extended processes and end-feet interacting with vessels (see arrowheads). (***H***) Vessels with positive-Casp3 labelling showed extravasation of albumin into the perivascular space and parenchyma **(*h*;** dotted line used to plot intensity profile). (***I***) Representative plot profile of mild albumin extravasation in FGR brain. All values are expressed as mean +/− SEM (minimum *n* = 6 for all groups). Two-way ANOVA with Tukey post-hoc test (* *p* < 0.05) (Scale bars: ***A*** - ***C***; 50μm, ***G;*** 20μm, ***H*** & ***h***; 10μm).

There was an increase in juxtavascular (*p* < 0.0001) and parenchymal (*p* = 0.002) Casp3^+^ labelling in FGR brains relative to NG (Fig. 8C&F). FGR+Ibu demonstrated a decrease in Casp3^+^ cells compared with untreated FGR (juxtavascular; *p* = 0.0005, parenchymal; *p* = 0.010). Further examination of Casp3^+^ cells found that these cells primarily co-localised with astrocytes (Fig. 8G). No co-localisation was observed with neurons or microglia at the vasculature. Vessels with juxtavascular Casp3 labelling were often associated with vessels displaying endogenous protein expression (Fig. 8H).

### Decreased expression of tight-junction protein Zonula Occluden-1 (ZO-1)

Tight junctions are a key component of the NVU and altered expression is associated with BBB disruption [81]. We examined the transmembrane tight junction proteins claudin-5 (Cldn5) and occluding (OCLN) and the cytosolic tight junction zonula occludens-1 (ZO-1). ZO-1 labelling showed a diffuse and disjointed pattern of labelling in FGR brain compared with the continuous labelling in all other groups (Fig. 9G). Co-localisation analysis showed a significant decrease in ZO-1 vessel coverage in FGR brain when compared with NG animals (*p* = 0.030), however no significant increase was observed following ibuprofen treatment (*p* = 0.990; Fig. 9H). Western blot analysis showed a significant decrease in ZO-1 protein expression in FGR compared with NG (*p* = 0.023), this decrease was not recovered following ibuprofen treatment (*p* =0.610; Fig. 9I). Robust labelling of Cldn5 was observed with strong co-localisation with endothelial cell marker (CD34) in all groups irrespective of pathology or treatment (*p* = 0.610; Fig. 9A&B); similarly western blot analysis found no difference in Cldn5 expression between any group (*p* = 0.124; Fig. 9C). OCLN displayed a more diffuse and lower intensity labelling pattern which was consistent across all groups (Fig. 9 D&E) with no significant differences in protein expression as determined by western blot analysis (*p* = 0.809; Fig. 9F).

**Fig. 9.**
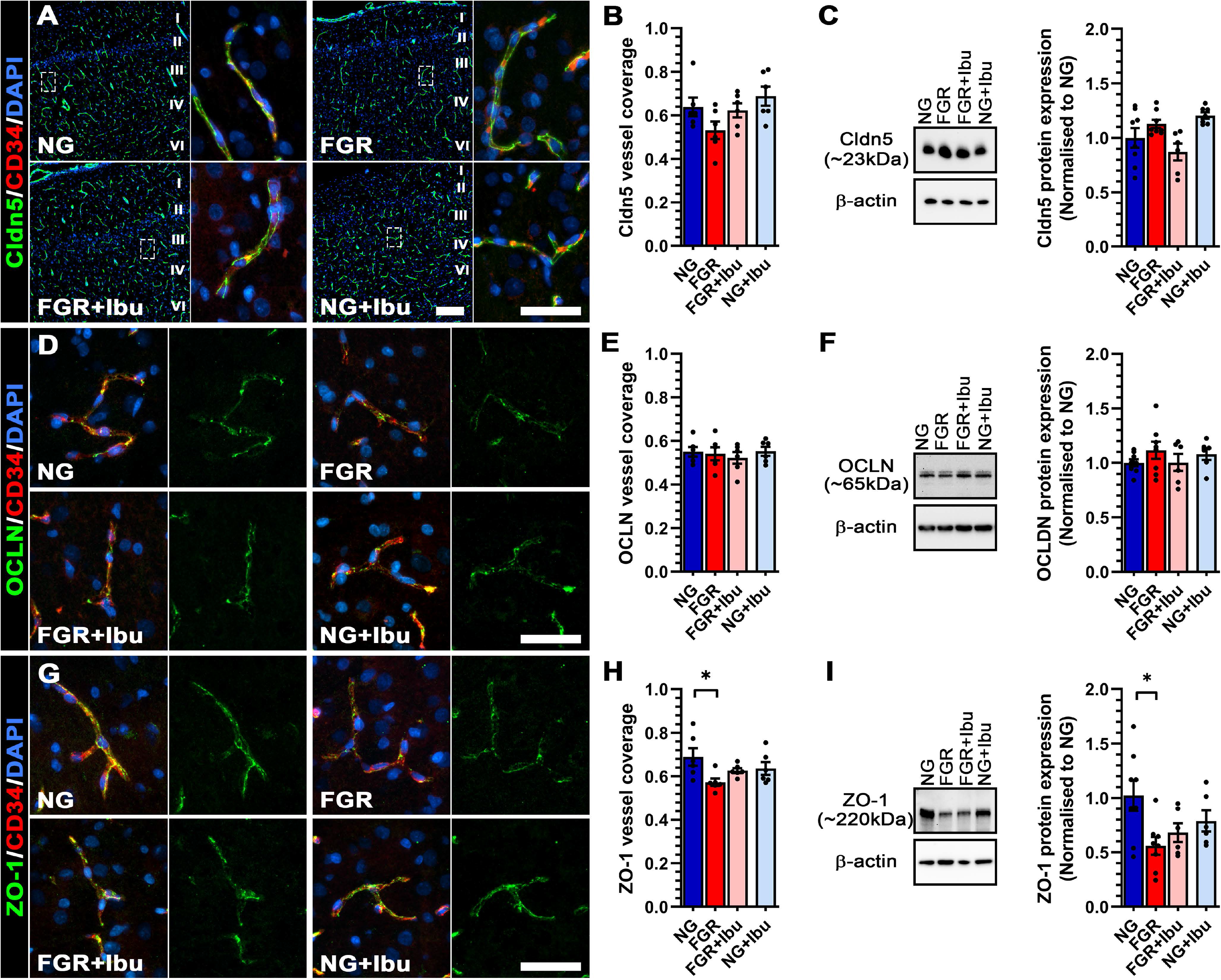
Decreased expression of tight junction protein ZO-1 in FGR brain. (***A***) Representative labelling of Claudin 5 (green; Cldn5) and endothelial cells (red; CD34) in the cortex at postnatal day 4. No significant difference in Cldn5 vessel coverage or protein expression was observed (***B*** *& **C*** respectively). (***D***) Tight junction protein Occludin (green; OCLN) showed low expression in all groups with more discrete labelling compared with other tight junction proteins investigated. (***E***) No significant alterations in OCLN vessel coverage or (***F***) protein expression was observed in any group. (***G***) Cytoplasmic tight junction protein Zonula Occludin-1 (green; ZO-1) demonstrated more diffuse labelling patterns in FGR compared with continuous patterns in other groups, which was associated with a subtle decrease in vessel coverage (***H***). (***I***) Western blot analysis showed a significant reduction in ZO-1 expression in FGR cortex compared with NG. All values are expressed as mean +/− SEM (minimum *n* = 6 for all groups). Two-way ANOVA with Tukey post-hoc test (\**p* < 0.05) (Scale bars: 50μm, low magnification scale bar in A: 200μm).

## Discussion

The NVU plays an essential role in progression of numerous CNS pathologies in the adult brain, such as Alzheimer’s disease (AD), multiple sclerosis (MS), stroke, and ischemia [83, 85]. The BBB in newborns is often considered immature, however a growing body of evidence suggests that while cellular function may differ from the adult, there is early establishment of functional and protective properties to meet the demands of the developing brain [77]. Here, we demonstrate clear evidence of NVU-alterations in the newborn FGR brain, with increased presence of endogenous proteins in the perivascular space and discrete-focal extravasation into the parenchyma. We further demonstrated reduced levels of ZO-1 (tight junction) protein and T-cell infiltration into the FGR brain. These alterations are attributed to the elevated pro-inflammatory profile in the FGR brain and the activated glial morphology reported here. Treatment with a common anti-inflammatory, ibuprofen, significantly decreased glial cell activation, ameliorated elevated cytokine levels, and resulted in normalised glial vessel interaction which was associated with reduced BBB-disruption. Our findings suggest that BBB-disruption in FGR may exacerbate early neuroinflammatory responses and therefore early targeting of inflammatory pathways in the clinical setting may provide a therapeutic approach in protecting the FGR newborn from adverse neurological outcomes.

### Mechanisms of action of ibuprofen on the NVU

Ibuprofen’s action on the NVU has not previously been examined in neonatal brain disorders. Previous studies investigating the neuroprotective potential of ibuprofen in compromised newborns have largely focused on neurons, white matter and inflammation in the brain parenchyma [14, 93, 94]. Ibuprofen, through inhibition of COX 1 and 2 activity, has various critical functions in the brain, through the reduction in the production of prostaglandins (PGs) and modulation of pro-inflammatory pathways. PGs and pro-inflammatory cytokines TNFα and IL-1β have major actions on blood vessels whereby TNFα and IL-1β affect vascular permeability and increase immune cell infiltration. The dosage of ibuprofen used in the current study significantly reduced the pro-inflammatory profile in the FGR brain as previously reported [94]. This reduction in inflammation in the brain microenvironment following ibuprofen treatment coincided with reduced disruption to the NVU composition and reduced infiltration of immune cells.

In a rat spinal cord injury model, ibuprofen significantly reduced lanthanum (an electron dense tracer) infiltration into endothelial cells and in the basal lamina, but extravasation of lanthanum in vesicles was mainly absent [78]. Ibuprofen also prevented the disturbances in the blood-spinal cord barrier permeability, edema formation, spinal cord blood flow changes, and cell reaction. In contrast, ibuprofen treatment amplified rather than decreased microvascular damage associated with seizures [75]. In this rat seizure study, ibuprofen administration increased the expression of TNFα and IL-1β in microvessels with aggravation of edema and microbleeds; however cerebral tissue was protected from inflammation following treatment [75].

### FGR newborn pig display altered vascular integrity and glial morphology

The neurovascular unit (NVU) is composed of neurons, glial cells (astrocytes, microglia, and oligodendroglia), vascular endothelial cells, and the basement membrane. The NVU plays a critical role in regulating cerebral homeostasis through maintenance of BBB integrity and cerebral blood flow [1, 85]. Studies of induced-FGR in the preterm and term lamb, through single umbilical artery ligation, indicate altered BBB-permeability which may be associated with brain pathology [15, 54]. Here we employed the use of MRI with gadolinium contrast agent to ascertain whether BBB-disruption could be identified *in vivo* in naturally occurring FGR piglets using a common clinical approach [64]. Studies in neurological disorders such as MS, which is characterised by inflammatory lesions of the white matter, have reported the formation of new lesions preceding BBB-disruption as detected by gadolinium enhancement [2]. However very few studies examine gadolinium for BBB-disruption in newborns. A neonatal stroke model showed minimal gadolinium enhancement 24h after reperfusion, even with microscopic evidence of BBB-disruption [27]. We detected *in vivo* alterations in MRI contrast enhancement in the FGR brain supporting the postmortem microscopic investigations of changes in vascular integrity. The T1 signal was increased (indicative of tissue gadolinium), while T2 was also increased (vascular gadolinium) even though vascular density was reduced. We observed a 30% decrease in collagen IV, the most abundant basement membrane protein, which provides support and adhesion for basal vascular endothelial cells (CD34) [89, 101] for which we report a 40% decrease in the parietal cortex of the FGR brain. The primary cells involved in production of collagen IV are endothelial cells, astrocytes, and pericytes [44, 84, 87, 91]. Thus, a loss or dysfunction of these cell types (discussed below) may contribute to loss of collagen IV. We also observed truncation and discontinuity of labelling of basement membrane and endothelial cells in the FGR brains with a significant reduction in proliferating cells. Studies by Castillo-Mendez *et al*., [15, 16] report a reduction in vascular density in the white matter of FGR lambs using the non-collagenous glycoprotein laminin as a marker of the basement membrane. Similarly, they also demonstrate a decrease in proliferating blood vessels in the brain at 24h postnatal age in the FGR lamb [16], supporting vascular disruption in the FGR brain.

### Inflammatory state alters glial morphology and interactions with brain microvessels in FGR brain

Glial cell changes have been extensively characterised following brain trauma. Changes in morphology and phenotype result in an altered profile of surface markers and expression of an array of inflammatory mediators [25, 39, 92, 94]. In FGR brain, we have shown that juxtavascular microglia and astrocytes demonstrate classical thickening and retraction of processes, and increased expression of pro-inflammatory mediators. Key upstream inflammatory initiators NF-κΒ, IL-1β and TNFα were expressed in juxtavascular microglia and astrocytes that interact with the NVU of FGR brains. These factors exhibit a multitude of innate and adaptive immune functions which are critical mediators in brain pathology and brain development [24, 58, 60]. Excess production of these inflammatory cytokines in both acute and chronic environments is associated with disease pathogenesis and brain dysfunction [19]. Microglia display dynamic motility behaviours following tissue damage, and preferentially associate with microvessels [29, 63]. Juxtavascular microglia have reported to be key players in BBB disruption following stroke [38], where inhibition of microglial activation after stroke promotes BBB integrity and viability [99]. A recent induced-systemic inflammation mouse study reported microglia have dual effects on BBB integrity [31]. Microglia have been shown to respond to inflammation by migrating towards and accumulating around cerebral blood vessels before any detectable changes in BBB permeability are evident. The initial juxtavascular microglial contact with endothelial cells protected the integrity of the BBB; however, as inflammation progressed, a more activated microglial phenotype dominated, resulting in phagocytosis of astrocytic end-feet and impairment of the BBB [31]. A mouse stroke study suggests phagocytic microglia may disrupt the BBB by engulfing endothelial cells [38]. Microglia were activated soon after reperfusion with expansion of their cellular processes towards blood vessels. All activated microglia in the damaged region were associated with, and engulfed cerebral blood vessels within 24h and concurrently broke down the BBB [38]. We observed an increase in activated juxtavascular microglia, hypertrophic microglia appeared to engulf some blood vessels in the FGR brain. The activated microglia were also in close contact with astrocytic end-feet and therefore may be acting in a phagocytic manner towards these and other cell types.

Utilising 3D reconstruction of electron microscopic images, Mathiisen *et al*., (2010) demonstrated almost complete ensheathment of brain microvessels by astrocytic end-feet under non-pathological conditions [57]. The interaction of astrocyte end-feet and brain microvessels is a key component in maintaining BBB-integrity, with studies demonstrating that under pathological conditions end-feet retract from vessels, exacerbating injury and altering BBB-permeability [36, 37, 96]. In the FGR brain, we report activation of juxtavascular astrocytes was associated with decreased end-feet ensheathment of the vasculature and may be a key mediator of BBB permeability. FGR sheep studies concur with this alteration to BBB, reporting a decrease in astrocyte end-feet coverage of blood vessels within white matter regions 24h after preterm and term birth [15, 54]. Further, in our current study, the astrocytes in contact with blood vessels in the FGR brain displayed enlarged end-feet indicative of change in functionality. These astrocytes may be no longer functioning as a support cell, rather are in a phenotypic reactive state with the ability to release inflammatory cytokines.

Activation of microglia and astrocytes results in the initiation of numerous inflammatory pathways in the order to resolve tissue damage. However, sustained release of cytokines and chemokines can exacerbate the inflammatory cycle contributing to NVU instability. The pro-inflammatory cytokines IL-1β and TNFα are well established as regulators of neuroinflammation but there is evidence to support them as key inducers of BBB dysfunction [86]. Data from several studies suggest microglial derived TNFα and IL-1β play a major role in BBB disruption [18, 22, 45]. Microglia activated in response to epileptic seizures release IL-1β which has been shown to downregulate the tight junction protein Cldn5 in endothelial cells resulting in disruption to BBB function [22]. TNFα binds to endothelial cells, leading to a downregulation of tight junctions and subsequent increase in BBB permeability in many central and peripheral inflammatory conditions [67, 97]. Microglial secreted TNFα has also been recognised as a key cytokine leading to endothelial cell programmed necrosis after stroke [18] [42]. These findings support our observations of TNFα and IL-1β expression in juxtavascular microglia in FGR brains and corresponding alterations to BBB permeability. CXCL10 is expressed in multiple CNS pathologies and strongly linked with alterations in BBB-permeability, although the mechanism through which it acts is unclear [6, 17, 90]. Further, in the current study, juxtavascular astrocytes showed strong TNFα expression in FGR brains. We also observed robust labelling of NF-κΒ in both astrocytes and microglia in the FGR newborn brain while only moderate labelling was noted in the other groups investigated. This prototypical pro-inflammatory pathway is likely triggered by the increased expression of IL-1β and TNFα [47, 49]. NF-κB may be essential in promoting microglial activation and TNFα release, which mediates endothelial programmed necrosis and accelerates BBB disruption after stroke injury [18]. Activation of this pathway may exacerbate and prolong the pro-inflammatory profile in the FGR brain with studies demonstrating inhibition of NF-κΒ signaling is associated with reduced inflammation and brain injury in rodent models of neuropathology [52, 73, 98].

Administration of ibuprofen significantly reduces parenchymal glial activation in FGR newborns [94]. In our current study, ibuprofen treated FGR animals showed a reduction in the number of activated juxtavascular microglia as well as parenchymal microglia with a return to the glial morphology observed in normally grown animals. Juxtavascular astrocytes in the treated FGR brains also returned to a resting morphology that resulted in more consistent end-feet coverage of microvessels. These findings coincided with a reduction in several inflammatory cytokines. This positive effect on the glial cells of the NVU in the FGR brain has also been shown with other anti-inflammatory treatments. Maternal melatonin treatment for the FGR fetal sheep resulted in improved astrocytic end-feet coverage of blood vessels and reduced BBB disruption [16]. It is therefore plausible that reducing the ongoing proinflammatory cycle in the FGR brain could allow the brain to function in a healthy way whereby the astrocytes and microglia provide protection and support to the BBB rather than disruption to the BBB.

The use of endogenous proteins (IgG and albumin) are well-established for detection of altered BBB-permeability [28, 66, 103]. A clear limitation of this approach is that they cannot be used to determine progression or duration of the ‘leak’ as these proteins readily diffuse once in the brain parenchyma [30]. The increase in perivascular albumin labelling we demonstrate in FGR animals may explain the observed increase in T2 vascular gadolinium. This MRI technique is unable to distinguish between vascular and perivascular concentrations of gadolinium, therefore the T2 concentrations we report may largely be confined to the perivascular space. Regions displaying BBB-disruption were associated with glial alterations as described above. We also observed neuronal and glial uptake of endogenous proteins in the FGR brain. In models of hypoxia-ischemia, neuronal uptake of IgG was associated with neuronal degeneration and reported to result in perturbed neuronal function and survival, and therefore may contribute to the progression of neuropathology [28, 61].

In the normally grown and ibuprofen treated FGR piglets, a reduction in the presence of perivascular albumin was observed, along with infrequent focal IgG extravasation in the cortex and white matter with strong astrocytic end-feet interaction around vessels. Further, the astrocytic processes appear to form a boundary around the focal leaks, suggesting containment of the leak or support of the vessels. This may be due to the reduced pro-inflammatory state of the brain whereby the astrocytes are able to function as a supportive cell rather than as a reactive phenotype. In agreement with our findings, melatonin administration to the FGR lamb reduced albumin extravasation in the brain and conferred protection on the BBB [16]. Umbilical cord blood-derived stem cell treatment to the FGR lamb also resulted in decreased albumin extravasation in brain when assessed at 24h postnatal age, with a concomitant reduction in brain injury [55]. Whether this protection to the BBB is directly via an anti-inflammatory effect of treatment is unknown. However, mouse studies suggest this may indeed be plausible [31, 99]. Minocycline treatment in a mouse model of systemic inflammation maintained BBB integrity by directly inhibiting microglial activation [31]. Further, inhibition of microglia in a mouse stroke functional microglia knockout model not only reduced extravasation of MRI contrast agent in the brain parenchyma but also resulted in a reduction in lesion size [38].

### BBB-permeability is associated with infiltration of peripheral immune cells

To our knowledge, this is the first report of T-cell infiltration into the newborn FGR brain. T-cell infiltration has been reported to promote neuroinflammation and cognitive decline in animal and human neuropathological studies [48, 59]. Mild suppression of immune responses in the FGR pig at postnatal day 24 have been demonstrated with a decrease in peripheral lymphocytes, more specifically CD3^+^CD4^+^ T-cells [5]. Combined with our findings of early CD3^+^ cell infiltration into the brain parenchyma, FGR newborns likely present a compromised immune response. Characterisation of changes in both peripheral and CNS immune cell populations is necessary to determine the contribution of immune dysfunction to the altered brain pathology we and others have reported in FGR newborns. We have not yet identified whether these T-cells belong to cytotoxic (CD8^+^) or T-helper (CD4^+^) populations, which would enable a better understanding of whether the CD3^+^ cell infiltration is a rescue phenotype or a population that contributes to neuropathology in the FGR brain.

The pro-inflammatory profile observed in FGR brain included expression of a number of chemokines and cytokines involved with chemotaxis of myeloid and lymphoid cells [4] in juxtavascular glia. Studies have reported juxtavascular astrogliosis coincides with alterations in BBB integrity, and suggest that BBB disruption precedes overt CNS infiltration by immune cells [3]. These studies propose that immune cell infiltration may therefore be a direct consequence of glial activation [3]. There is evidence of activated juxtavascular microglia accumulating in sites of lymphocyte infiltration in a bacterial endotoxin lipopolysaccharide postnatal rat model [95]. Activated microglia are known to release chemoattractants for lymphocytes in addition to pro-inflammatory mediators that stimulate astrocyte activation [4]. We observed elevated expression of CCL2 (MCP-1) which is known to contribute to site-specific infiltration of lymphocytes into the brain [13]. These findings demonstrate the interplay between glial activation, NVU dysfunction and the drive to promote recruitment of peripheral infiltrates, which if left unchecked may result in sustained and deleterious effects on the newborn FGR brain.

We observed labelling of the tight junction protein Cldn-1 in the FGR brain, with less labelling detected in the cortex and white matter in the normal brain [7]. Claudin-1 expression was co-localised with GFAP-positive astrocytes throughout the parenchyma as well as in juxtavascular astrocytes. The pro-inflammatory cytokine IL-1β has been shown to induce astrocytic expression of Cldn-1 which acts to corral activated T-cells in the brain [33]. Activated T-cells in turn release metalloproteinases (MMPs) 3, 7, and 9, to degrade Cldn-1 protein. The authors propose a subtle ongoing struggle between activated astrocytes and T-cells attempting infiltration of the brain parenchyma. The expression of Cldn-1 in astrocytes may therefore be a compensatory mechanism to provide a ‘secondary’ barrier to infiltration of peripheral immune cells into the CNS. By dampening the pro-inflammatory environment, with ibuprofen, we propose that glial cells return to their resting morphological state, allowing normal function and reduced energy demands. This normalised glial interaction at the NVU provides a ‘tighter’ more functional BBB, which minimises leak of serum proteins and peripheral cell infiltration into the brain. Together these ameliorated changes provide the newborn brain a healthier environment for early postnatal brain development, which in turn is likely to improve long-term neurodevelopmental outcomes.

### Ibuprofen treatment reduces apoptosis in the brain parenchyma and at the NVU

There was an evident increase in the number of Casp3^+^/GFAP^+^ juxtavascular astrocytes in the FGR brain. A study in human AD patients showed the presence of degenerating astrocytes as labelled with an antibody specific to GFAP-caspase cleavage product [65]. These degenerating astrocytes co-localised with caspase-3 antibodies and were found in proximity to the neurovasculature. Further, a significantly higher number of caspase-3 positive cells are observed on GLUT-1 positive endothelial cells in FGR lambs with a concurrent decrease in the astrocytic end-feet coverage of blood vessels [16]. Together with our observations, these findings suggest activation of caspase-3 pathways and cleavage of GFAP (cytoskeletal proteins) may contribute to decreased astrocyte end-feet interaction with brain microvessels and subsequently result in BBB-permeability. Following ibuprofen treatment, we observed a significant decrease in Casp3-positive cells in the FGR brain. Ibuprofen has been shown to have a direct effect on apoptosis [80]; therefore the positive impact of ibuprofen treatment on the NVU may not only be due to its anti-inflammatory effect, but also its anti-apoptotic actions. A similar effect has been shown following melatonin treatment in the FGR lamb. Maternal melatonin treatment significantly reduced the number of caspase-3 positive cells associated with blood vessels in the white matter of FGR lamb brains [16]. It is plausible that these treatments provide neuroprotection through multiple actions that target apoptosis and inflammation in the FGR brain.

### Decreased expression of tight junction protein ZO-1 in FGR brain

Tight junction proteins are key regulators of BBB-integrity, with numerous studies demonstrating that loss or mutations in TJs are associated with BBB-disruption. Depletion of Cldn-5 induces BBB-disruption in knockout mice [68] and in cultured human brain endothelial cells [53], while OCLN knockout resulted in growth retardation but no significant alterations in BBB-function [76]. Our investigations of these key TJ proteins found no change in Cldn-5 and OCLN, however the cytosolic TJ protein, ZO-1 was significantly decreased in FGR brain. The absence of disruption to Cldn-5 and OCLN protein levels may explain why overt BBB-disruption was not observed in FGR brains [32, 100]. However, in neonatal HI piglet with extensive brain injury and BBB disruption (IgG extravasation), no changes were observed at the mRNA level of these TJs, and only loss of ZO1 and OCLN protein in the parietal cortex of HI animals with seizures [28].

While expression of TJs were generally maintained in the FGR brain, interactions between these key regulatory proteins may be affected. Under pro-inflammatory endothelial conditions, Cldn-1 is expressed at the endothelial cells and reported to disrupt the interaction between Cldn-5 and ZO-1 which in turn reduces BBB-integrity [79]. While we report relatively normal expression levels of TJ proteins in FGR, it is plausible that Cldn-1 expression at the microvasculature may contribute to altered TJ protein interactions and subsequently result in reduced BBB-integrity.

It is also possible the TJ proteins may have undergone redistribution. Models of stroke have demonstrated movement of ZO-1 from the membrane to cytoplasm during reperfusion [102]. ZO-1 loss and TJ redistribution in traumatic brain injury has also been shown to be associated with IL-1β [69]. As we have demonstrated a significant increase in IL-1β, but also a loss in ZO-1, redistribution of TJs in the FGR brain cannot be discounted. The present study used whole protein preparations and did not examine the subcellular fractions of brain tissue; further investigation is required to ascertain whether TJ proteins are significantly redistributed in FGR brains. Studies demonstrating significant alterations in TJ protein expression generally show overt BBB-disruption [102]. Given the mild BBB-disruption reported in the present study and relatively normal expression of total protein for each TJ, our findings would indicate they are not the key mediator involved in the disruption observed. These findings indicate only subtle structural alterations to the vasculature and TJs of FGR brain suggestive that the BBB-permeability observed is likely due to altered function or interaction rather than loss of NVU components.

## Conclusion

FGR is associated with glial activation which results in alterations to juxtavascular astrocytes of the NVU. Altered glial-interaction contributes to BBB-permeability and leakage of peripheral proteins and infiltrates into the brain parenchyma, potentially further exacerbating the inflammatory environment. Ibuprofen treatment reduced neuroinflammatory glial responses and was associated with improved NVU interactions. As ibuprofen crosses the BBB [72], it is likely it may act centrally and systemically to modulate inflammation. As there is a cyclic inflammatory event in the FGR brain, reducing inflammation either peripherally or centrally would likely be beneficial. However, further research into the direct interactions of ibuprofen on the NVU, as well as determining long-term efficacy of this treatment are warranted.

## Declarations

### Ethics approval

The University of Queensland granted Ethics approval for this study and the study was carried out in accordance with the National Health and Medical Research Council guidelines (Australia) and ARRIVE guidelines.

### Author contributions

Kirat Chand was involved in obtaining funding and responsible for all laboratory aspects of the project, data analysis, interpretation, writing and editing the manuscript. Stephanie Miller was involved in conducting animal experiments and editing the manuscript. Gary Cowin was involved in MRI acquisition, data analysis and editing the manuscript. Lipsa Mohanty undertook all western blotting and edited the manuscript. Paul Colditz was involved in obtaining funding, critical revision, and editing the manuscript. Stella Tracey Bjorkman was involved in obtaining funding, critical revision, and editing the manuscript. Julie Wixey was involved in obtaining funding, study conception and design, conducting animal experiments, acquiring data, critical revision, and writing the manuscript.

## Acknowledgments

The authors acknowledge the facilities and scientific and technical assistance of the National Imaging Facility, a National Collaborative Research Infrastructure Strategy (NCRIS) capability, at the Centre for Advanced Imaging, University of Queensland. The authors would also like to thank Lara Jones, Elliot Teo, and John Luff for assistance with animal experimentation.

Financial support was provided by The National Health and Medical Research Council (Australia) (Grant ID: APP1147545), Royal Brisbane and Women’s Hospital Foundation, and Children’s Hospital Foundation (Grant ID 50217) grants. The authors acknowledge the facilities and scientific and technical assistance of the National Imaging Facility, a National Collaborative Research Infrastructure Strategy (NCRIS) capability, at the Centre for Advanced Imaging, University of Queensland. The authors would also like to thank Lara Jones, Elliot Teo, and John Luff for assistance with animal experimentation.

---

**Supplementary Fig. 1.**
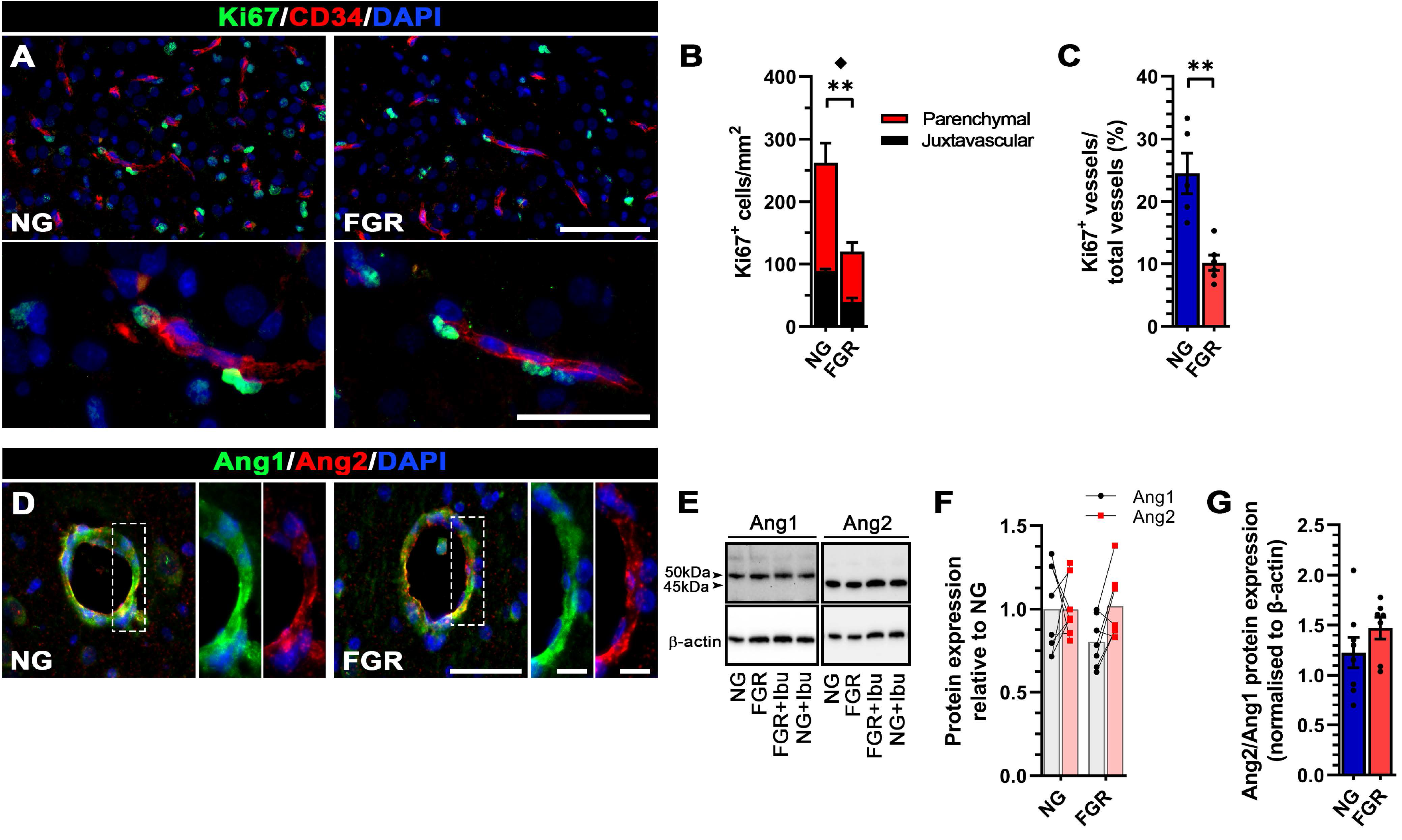
Reduced proliferation and angiogenesis was observed in FGR. (***A***) Representative labelling of Ki67 (green) and endothelial cells (red; CD34) in cortex demonstrating an overall decrease in proliferating cells in the FGR brain. (***B***) Significant reductions in Ki67^+^ cell counts were found in the parenchyma and juxtavascular regions. (***C***) Analysis of Ki67^+^ vessels relative to the total number of vessels demonstrated a significant reduction in vessels undergoing proliferation in the FGR group. (***D***) Examination of angiogenic markers angiopoietin-1 (Ang 1; green) & angiopoietin-2 (Ang2; red) found similar labelling patterns in both NG and FGR. (***E*** *& **F***) Western blot analysis showed no significant overall difference in expression between NG and FGR. (***G***) Analysis of Ang2/Ang1 ratio demonstrated a mild but non-significant increase in FGR brains. All values are expressed as mean +/− SEM (minimum *n* = 6 for NG and FGR). Unpaired student’s t-test (^*^*p* < 0.05 (perivascular); \*\**p* < 0.01 (parenchymal)) (Scale bars: 50μm; ***D*** high magnification: 10μm).

**Supplementary Fig. 2.**
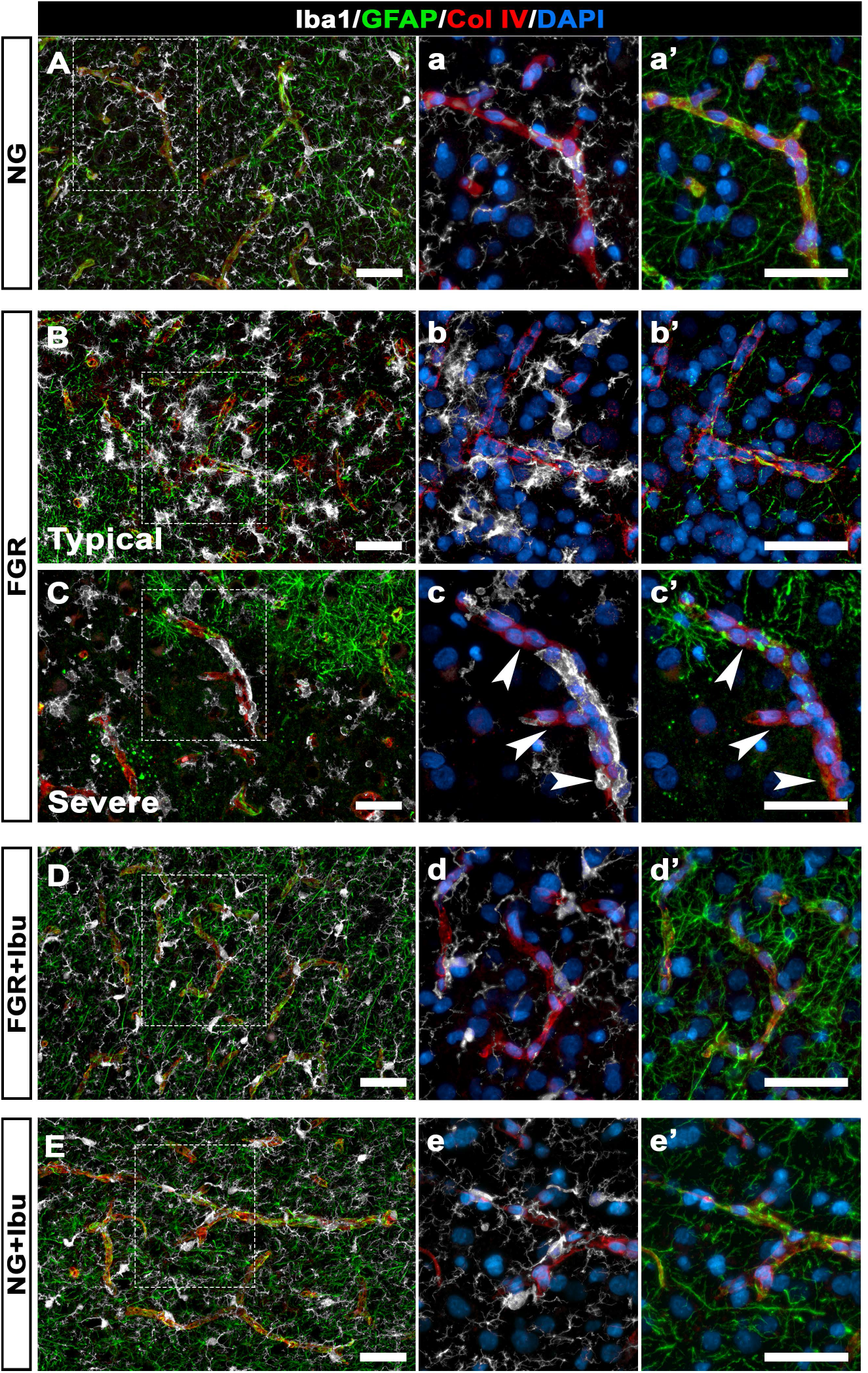
Glial interactions with transverse vessels demonstrate increased microglia and decreased astrocytic interaction in FGR. Representative labelling of microglia (grey) and astrocytes (green) interactions with vasculature (red) of the neonate brain. (***A***) NG demonstrated resting glial morphology, with long thin process extensions forming an organised network of tight interactions along the vasculature. (***a***) Ramified microglia are present in close proximity, while astrocytes showed end-feet interaction with vasculature (***a’***). (***B***) Representative labelling for FGR demonstrated an increase in activation of glia based on morphology, with retraction of extended processes and dense cellular bodies. FGR displayed close interaction of activated microglia with the vessel (higher magnification shown in ***b***). (***b’***) There was an evident decrease in GFAP labelling spanning the length of vessel. (***C***) In severe instances there was a high degree of activated juxtavascular microglia (***c***) and significant loss GFAP labelling along the vasculature with some regions completely void of astrocyte interaction (arrowheads; ***c’***). (***D*** & ***E***) Ibuprofen treated FGR and NG groups displayed glial morphology comparable with untreated NG, indicating a normalised resting state glial morphology and strong interaction with the vasculature (***d-d’*** & ***e-e’***). (Scale bars: 50μm).

**Supplementary Fig. 3.**
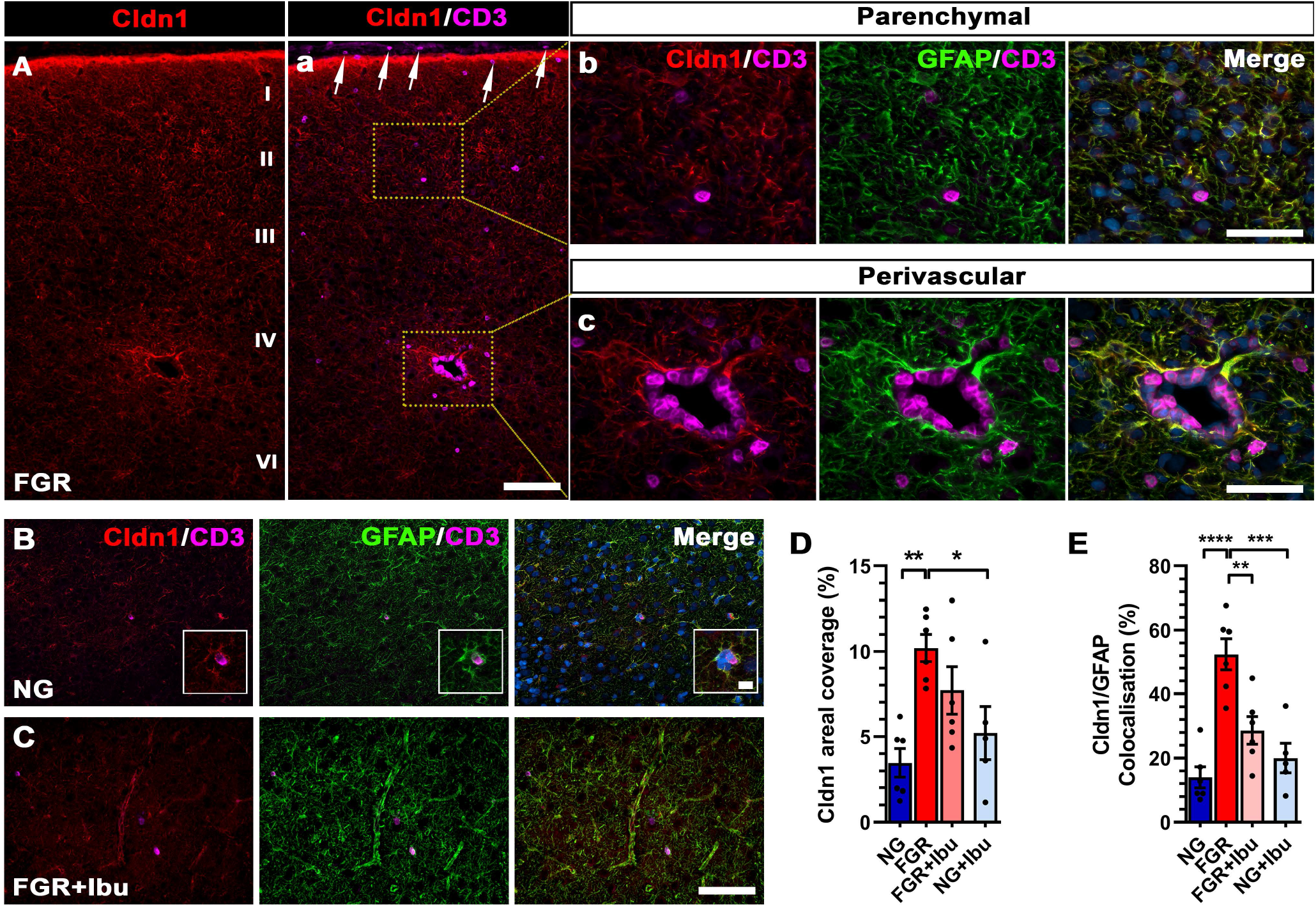
Elevated Claudin-1 labelling in FGR brain. (***A***) Robust labelling of tight junction protein claudin-1 (Cldn1; red) was observed at the pial surface, across cortical layers, and white matter of FGR brains at postnatal day 4. CD3^+^ cells were observed in regions with high degree of Cldn1 labelling (***Aa***). Cldn1 labelling showed strong co-localisation to astrocytes (GFAP; green) which displayed activation as characterized by disorganized and thickened process morphology at both parenchymal and perivascular regions (***Ab*** & ***Ac***). NG and FGR+Ibu brains showed limited Cldn1 labelling, predominantly restricted to astrocyte end-feet interacting with CD3^+^ cells and the vasculature (***B*** & ***C*** respectively). (***D***) Quantification of Cldn1 areal coverage demonstrated a significant elevation in FGR, which was partially ameliorated following ibuprofen administration. (***E***) Astrocytes in FGR brains displayed significantly elevated expression of Cldn1 compared with all groups. Ibuprofen administration significantly reduced the degree of Cldn1 labelling in FGR animals. All values are expressed as mean +/− SEM (minimum *n* = 6 for all groups). Two-way ANOVA with Tukey post-hoc test (\**p* < 0.05; \*\**p* < 0.01; \*\*\**p* < 0.001; \*\*\*\**p* < 0.0001) (Scale bars: 100μm, ***Ab*** & ***c***: 50μm, ***B*** & ***C*** inserts: 10μm).

**Supplementary Table 1:**

| **Primary Antibody** | **Host** | **Dilution** | **Catalogue#** |
| --- | --- | --- | --- |
| Anti-Col IV | Rabbit | 1:1000 | Abcam (ab6586) |
| Anti-CD34 | Goat | 1:1000 | R&D Systems (AF3890) |
| Anti-Ki67 | Rabbit | 1:500 | Abcam (ab15580) |
| Anti-Ang1 | Rabbit | 1:500 | Abcam (ab8451) |
| Anti-Ang2 | Rabbit | 1:200 | Abcam (ab153934) |
| Anti-Iba-1 | Goat | 1:1000 | Abcam (ab5076) |
| Anti-GFAP | Mouse | 1:1000 | Sigma-Aldrich (G3893) |
| Anti-NF-κB | Rabbit | 1:1000 | Abcam (ab16502) |
| Anti-TNF-α | Rabbit | 1:500 | Abcam (ab183218) |
| Anti-IL-1b | Rabbit | 1:500 | Abcam (ab9722) |
| Anti-IL4 | Rabbit | 1:500 | Abcam (ab9622) |
| Anti-CXCL10 | Rabbit | 1:500 | Abcam (ab9807) |
| Anti-CCL2 | Rabbit | 1:200 | Invitrogen (PA5-34505) |
| Anti-CCL3 | Rabbit | 1:500 | Invitrogen (PA5-86025) |
| Anti-porcine IgG | Goat | 1:1000 | JIR (114-005-003) |
| Anti-Albumin | Sheep | 1:1000 | Abcam (ab8940) |
| Anti-CD3 | Rat | 1:1000 | Abcam (ab11089) |
| Anti-Claudin 1 | Rabbit | 1:1000 | Abcam (ab15098) |
| Anti-Cleaved Caspase-3 | Rabbit | 1:500 | Cell Signaling (#9661) |
| Anti-αSMA | Mouse | 1:1000 | Abcam (ab7817) |
| Anti-Cldn5 | Rabbit | 1:1000 | Abcam (ab131259) |
| Anti-OCLN | Rabbit | 1:1000 | Abcam (ab222691) |
| Anti-ZO1 | Rabbit | 1:1000 | Abcam (ab96587) |
| **Secondary Antibody** |  |  |  |
| α-Rat Alexafluor 488 | Donkey | 1:1000 | Invitrogen (A-21208) |
| α-Rabbit Alexafluor 488 | Donkey | 1:1000 | Invitrogen (A-21206) |
| α-Goat Alexafluor 568 | Donkey | 1:1000 | Invitrogen (A-11057) |
| α-Mouse Alexafluor 568 | Donkey | 1:1000 | Invitrogen (A10037) |
| α-Rabbit Alexafluor 594 | Donkey | 1:1000 | Invitrogen (A-21207) |
| α-Goat Alexafluor 647 | Donkey | 1:1000 | Invitrogen (A-21447) |
| α-Mouse Alexafluor 647 | Donkey | 1:1000 | Invitrogen (A-31571) |
| DAPI (4',6-Diamidino-2-Phenylindole, Dihydrochloride) | - | 1:2000 | Invitrogen (D1306) |

